# Genomic biosurveillance of the kiwifruit pathogen Pseudomonas syringae pv. actinidiae biovar 3 reveals adaptation to selective pressures in New Zealand orchards

**DOI:** 10.1101/2024.10.21.616179

**Authors:** Lauren M. Hemara, Stephen M. Hoyte, Saadiah Arshed, Magan M. Schipper, Peter Wood, Sergio L. Marshall, Mark T. Andersen, Haileigh R. Patterson, Joel L. Vanneste, Linda Peacock, Jay Jayaraman, Matthew D. Templeton

**Affiliations:** School of Biological Sciences, The University of Auckland, Auckland, NZ; The New Zealand Institute for Plant and Food Research Limited, Mount Albert Research Centre, Auckland, NZ; The New Zealand Institute for Plant and Food Research Limited, Ruakura, NZ; The New Zealand Institute for Plant and Food Research Limited, Hawke’s Bay, NZ; Kiwifruit Vine Health, Mount Maunganui, NZ; Bioprotection Aotearoa, Lincoln University, Lincoln, New Zealand

## Abstract

In the late 2000s, a pandemic of *Pseudomonas syringae* pv. *actinidiae* biovar 3 (Psa3) devastated kiwifruit orchards growing susceptible yellow-fleshed cultivars. New Zealand’s kiwifruit industry has since recovered, following the deployment of the tolerant cultivar ‘Zesy002’. However, little is known about the extent to which the Psa population is evolving since its arrival. Over 500 Psa3 isolates from New Zealand kiwifruit orchards were sequenced between 2010 and 2022, from commercial monocultures and diverse germplasm collections. While effector loss was previously observed on Psa-resistant germplasm vines, effector loss appears to be rare in commercial orchards, where the dominant cultivars lack Psa resistance. However, a new Psa3 variant, which has lost the effector *hopF1c*, has arisen. The loss of *hopF1c* appears to have been mediated by the movement of integrative conjugative elements introducing copper resistance into this population. Following this variant’s identification, *in planta* pathogenicity and competitive fitness assays were performed to better understand the risk and likelihood of its spread. While *hopF1c* loss variants had similar *in planta* growth to wild-type Psa3, a lab-generated Δ*hopF1c* strain could outcompete wild-type on select hosts. Further surveillance was conducted in commercial orchards where these variants were originally isolated, with 6.6% of surveyed isolates identified as *hopF1c* loss variants. These findings suggest that the spread of these variants is currently limited, and they are unlikely to cause more severe symptoms than the current population. Ongoing genome biosurveillance of New Zealand’s Psa3 population is recommended to enable early detection and management of variants of interest.

## Introduction

Kiwifruit (*Actinidia* spp.) is a valuable perennial crop threatened by the bacterial pathogen *Pseudomonas syringae* pv. *actinidiae* (Psa). Psa biovar 3 (Psa3) spread throughout kiwifruit-growing regions worldwide during a pandemic in the late 2000s, causing significant economic losses (McCann et al., 2013; Vanneste, 2017). In 2010, an incursion of Psa3 was discovered in Te Puke, New Zealand’s main kiwifruit growing region (Everett et al., 2011). The following year, Psa3 devastated kiwifruit orchards growing susceptible *Actinidia chinensis* var. *chinensis* cultivars (Everett et al., 2011). Currently, Psa3 is the only biovar in New Zealand, alongside the closely related *Pseudomonas syringae* pv. *actinidifoliorum* (Pfm) (Vanneste et al., 2013; Cunty et al., 2015). Replacing *A. chinensis* var. *chinensis* ‘Hort16A’ with the less susceptible cultivar ‘Zesy002’ has helped the kiwifruit industry recover from the impact of this bacterial canker disease (Vanneste, 2017). However, Psa remains a persistent challenge to growers, requiring significant time and expense to manage through chemical controls, biological controls, and orchard hygiene practices.

New Zealand’s Psa3 population has continued to evolve and adapt to the environmental challenges presented in kiwifruit orchards, as exemplified by the acquisition of copper resistance from the local microbiome (Colombi et al., 2017). Furthermore, several Psa isolates from *A. arguta* vines have lost recognised effectors, making these variants more virulent on a kiwiberry species, typically considered Psa-resistant (Hemara et al., 2022).

Effectors of host-adapted pathogens interact with host targets to aid pathogen entry, suppress host immunity, and extract nutrients, ultimately benefiting pathogen virulence and establishing disease (Bent & Mackey, 2007; Chisholm et al., 2006; Xin et al., 2018; Zipfel, 2009). However, these effectors can also be recognised by plant resistance proteins, with effector-triggered immunity producing a robust immune response, preventing the establishment of disease and conferring host resistance (Nomura et al., 2005; Chisholm et al., 2006; Mur et al., 2008; Ngou et al., 2021; Yuan et al., 2021). Pathogens, in response, may undergo effector gene gain, loss, mutation and pseudogenisation to counter evolving plant immune systems and maintain their pathogenicity. Therefore, monitoring effector gain and loss in New Zealand’s Psa3 population is of particular importance to capture potential changes in virulence and the ability of this population to overcome disease controls, including host genetics.

Little is known about the full extent to which New Zealand’s Psa3 population may be adapting to its hosts since its introduction into commercial kiwifruit orchards. Further still, the exact genetic and immune mechanisms underlying the Psa tolerance of cultivars like ‘Zesy002’ are unknown. Therefore, new Psa variants may emerge that overcome this tolerance, thereby threatening New Zealand’s kiwifruit industry once more. Furthermore, new cultivars continue to be developed and commercially released, including the new red-fleshed cultivar *A. chinensis* var. *chinensis* ‘Zes008’. Each monoculture, with its unique genetics, may exert different selection pressures on this Psa3 population. Genome biosurveillance, early variant detection and subsequent pathogenicity characterisation are critical to ensure we can respond appropriately to emerging adaptations in the Psa population. The decreasing cost of whole genome sequencing has allowed us to conduct an in-depth and longitudinal genome biosurveillance programme to understand how Psa responds to selection pressures in the orchard environment and to reconstruct the trajectory of its adaptation over the past decade. This paper reports on the longitudinal genome surveillance of a clonal lineage of Psa3 in New Zealand kiwifruit orchards from 2010 to 2022.

## Results

Between 2010 and 2022, 571 Psa3 isolates were collected and sequenced from kiwifruit-growing regions in New Zealand’s North Island (Fig. 1A, B). Most of these isolates were sampled between 2017 and 2022 as part of a concerted genome biosurveillance effort (Hemara et al., 2022, 2024; Hoyte et al., 2024). This built upon preliminary sequencing research conducted by Plant & Food Research, Massey University, and The University of Otago from 2010 to 2016, during the initial years of the Psa3 incursion (Butler et al., 2013; McCann et al., 2013; Templeton et al., 2015; McCann et al., 2017; Colombi et al., 2017; Straub et al., 2018; Poulter et al., 2018). Of these, 513 isolates originated from commercial *A. chinensis* orchards growing monocultures of *A. chinensis* var. *chinensis* ‘Zesy002’ and *A. chinensis* var. *deliciosa* ‘Hayward’ (Fig. 1C). A further 58 Psa isolates were sampled from diverse *Actinidia* germplasm vines across Plant & Food Research’s research orchards in the North Island (Fig. 1C; Hemara et al., 2022, 2024). The New Zealand Psa3 population has a star-shaped core single nucleotide polymorphism (SNP) phylogeny, indicative of rapid expansion from a clonal origin (Fig. 1C). It is largely accepted that New Zealand’s Psa3 population was founded by a near-clonal introduction, represented by Psa3 V-13 (McCann et al., 2013, 2017).

**Figure 1.**
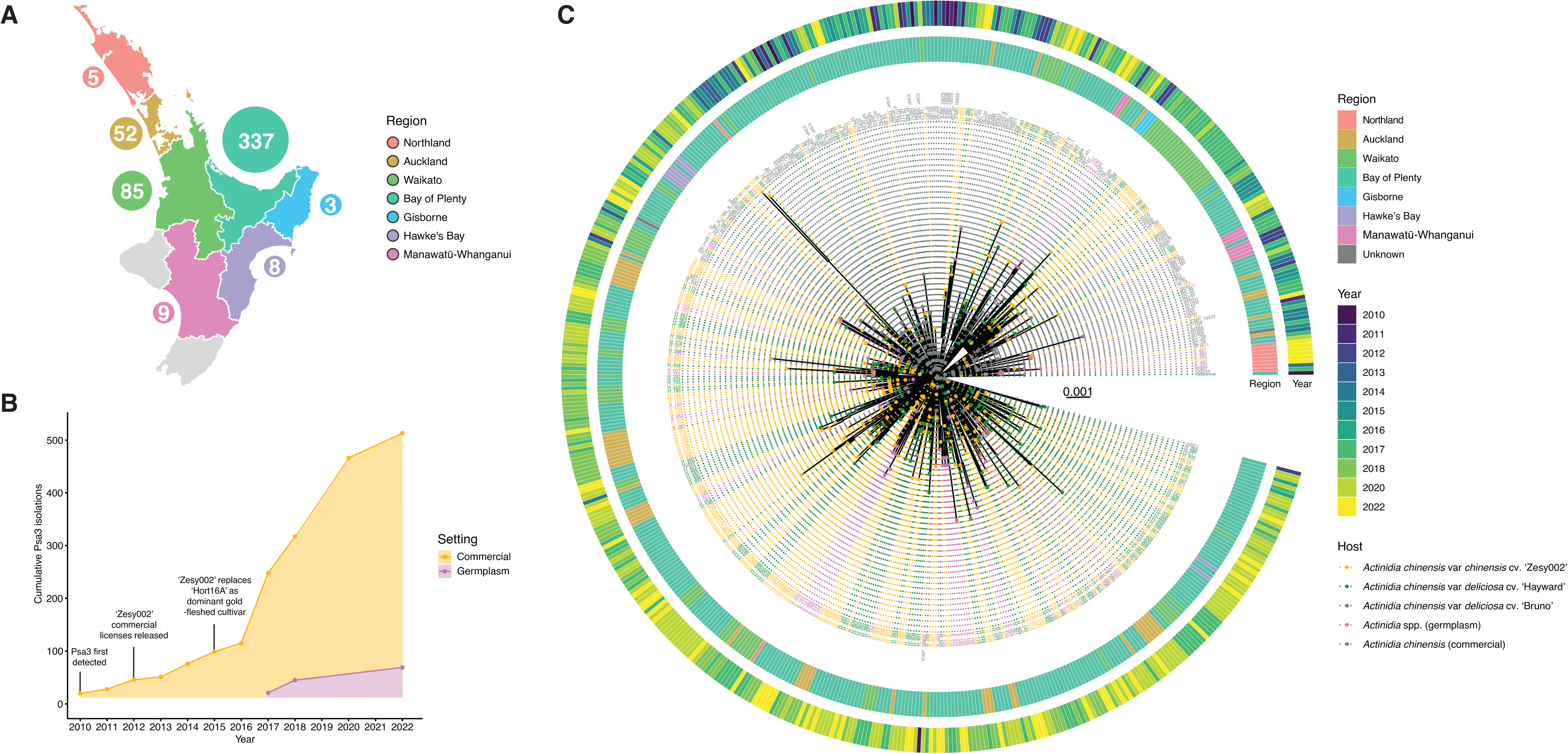
Psa3 isolates collected and sequenced from New Zealand kiwifruit orchards following the 2010 incursion. (A) Geographic distribution of Psa3 isolates collected from commercial kiwifruit orchards across the North Island of New Zealand. Bubble size is indicative only and does not directly correspond to the sample number. (B) Psa3 isolates cumulatively collected and sequenced from 2010 to 2022 from New Zealand kiwifruit orchards. Isolations from commercial *A. chinensis* cultivars are shown in orange; isolations from research orchard germplasm collections are shown in pink (Hemara et al., 2022, 2024). The x-axis represents the year of isolation for isolates that were later sequenced. (C) Core SNP phylogeny of New Zealand Psa3 isolates. A core SNP phylogeny of New Zealand Psa3 isolates was produced with Snippy (version 4.6.0) relative to the reference Psa3 V-13. Host and year of isolation are indicated.

Through comparison of these new isolates to the reference strain Psa3 V-13, isolated at the beginning of the pandemic (McCann et al., 2013), new variants were identified that have arisen through *de novo* mutation or the acquisition of novel genetic content through horizontal gene transfer (HGT). Psa3 V-13, which represents the original clonal population first introduced into New Zealand, carries an enolase-encoding, or *Tn6212*-carrying integrative conjugative element (ICE), which was predicted to be linked to virulence, likely to have a role in helping bacteria grow on preferred carbon sources *in planta* (Colombi et al., 2024). Given the widespread nature of copper spraying in kiwifruit orchards, it is perhaps unsurprising that copper resistance (Cu^R^) elements were the most common genomic acquisition, with over 65% of isolates carrying *copABCD* on an ICE or plasmid (Fig. S1). In Psa3, the acquisition of Cu^R^ ICE elements typically requires the excision and replacement of the *Tn6212-*encoding ICE. The most prevalent Cu^R^ element was PacICE10, carried by over 40% of isolates (Fig. S1). PacICE10 also conferred the highest copper resistance of all ICEs identified (Fig. S2). The clear dominance of PacICE10 over other elements suggests that this element has been strongly selected for.

Outside of ICE excision, gene deletions were less frequent than gene gain events. However, several elements appeared to have been deleted numerous times. The most frequent deletion was a 38 kb deletion excising a four-gene cassette *qatABCD* phage defence system (Fig. 2A; Gao et al., 2020; Lin et al., 2020), present in 34 commercial isolates and eight germplasm isolates. Another deletion of interest was that of a urease biosynthetic gene cluster (Fig. 2B), which had been lost in 23 *A. chinensis*-derived isolates and four germplasm-derived isolates. Interestingly, both of these deletions appear to have emerged independently over 20 times. Further, albeit rarer, deletions resulted in the loss of an achromobactin biosynthetic gene cluster (Fig. 2C) and an element carrying chemotaxis genes, including *cheY* and *pilT* (Fig. 2D). However, although many of these deletions occurred in multiple lineages (Fig. S3), the selective pressures driving gene loss remain unclear.

**Figure 2.**
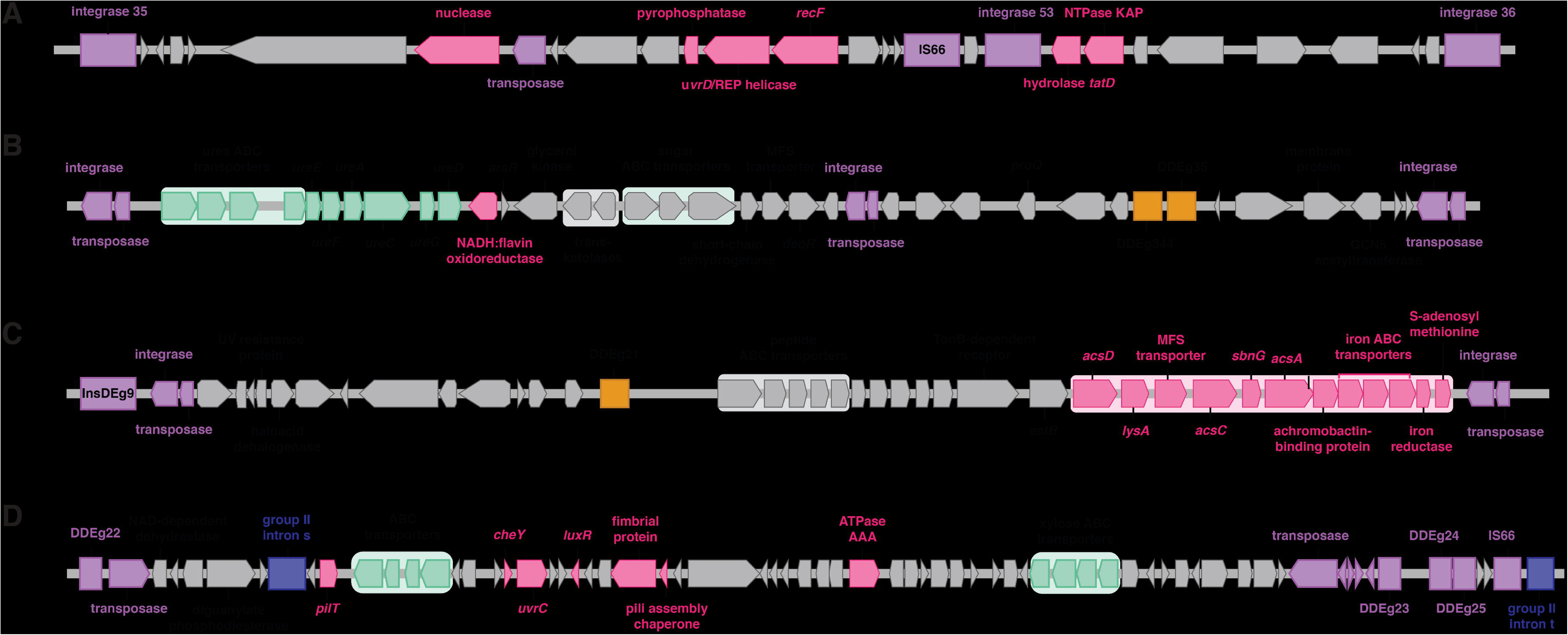
Gene deletions of interest in New Zealand’s Psa3 population. (A) Schematic of a 38 kb chromosomal gene deletion (6,474,001 – 6,512,000 bp), including nuclease, pyrophosphatase, *recF,* hydrolase *tatD,* and NTPase KAP. (B) Schematic of a 42 kb chromosomal gene deletion (3,740,001 – 3,782,000 bp) spanning the urease biosynthetic gene cluster. (C) Schematic of a 53.7 kb chromosomal gene deletion (2,923,241 – 2,976,955 bp) spanning the achromobactin biosynthetic gene cluster. (D) Schematic of a 66 kb chromosomal gene deletion (3,213,001 – 3,279,000 bp), including *pilT,* ABC transporters, *cheY, uvrC, luxR,* and pilus assembly proteins.

Alongside the acquisition of copper resistance (Colombi et al., 2017), one of the most significant changes observed to date in New Zealand’s Psa3 population has been effector loss on Psa-resistant germplasm vines (Hemara et al., 2022, 2024). Interestingly, no further effectors appear to have been gained by New Zealand’s Psa3 population since its introduction. On the other hand, effector loss appears to be highly host-dependent and differed significantly between isolates derived from germplasm collections and commercial orchards. Twenty percent of isolates collected from germplasm vines had lost one or more effectors (Hemara et al., 2022, 2024). In contrast, less than 1% of the commercial orchard Psa3 isolates surveyed in this study had lost an effector; these five Psa3 isolates all lost the same effector – *hopF1c* (Table 1). HopF1c is predicted to have ADP-ribosyl transferase function, supressing immune signalling cascades (Wang et al., 2010; Zhou et al., 2014). Interestingly, HopF1c appears to be recognised across both resistant and susceptible kiwifruit hosts (Hemara et al., 2022; Jayaraman et al., 2023). The *hopF1c* excision appears to have emerged three separate times in independent lineages from three spatially distinct orchards (Table 1; Fig. S3). Orchards A and B are approximately 9 km apart in the Bay of Plenty, while Orchard C is nearly 200 km south in the Hawke’s Bay region (Table 1).

**Table 1.** New Zealand Psa3 isolates lacking hopF1c. All isolates are from commercial *Actinidia chinensis* orchards.

| Isolate | Year | Cultivar | Orchard | Region |
| --- | --- | --- | --- | --- |
| G_121 | 2018 | ‘Zesy002’ | A | Bay of Plenty |
| H_407 | 2020 | ‘Hayward’ | B | Bay of Plenty |
| B_439 | 2020 | ‘Bruno’ | C | Hawke’s Bay |
| B_440 | 2020 | ‘Bruno’ | C | Hawke’s Bay |
| G_441 | 2020 | ‘Zesy002’ | C | Hawke’s Bay |

Following identification through comparative genomics, the loss of *hopF1c* was confirmed by multiplex-PCR using Psa ITS and *hopF1c*-specific primers (Fig. S4, S5, S6). In all *hopF1c* deletion isolates, the deletion of *hopF1c* was accompanied by the excision of the downstream*Tn6212*-carrying ICE and replacement with a Cu^R^-encoding ICE. Short and long read sequencing indicated that Psa3 G_121, the first *hopF1c* loss isolate identified, has a 93 kb deletion spanning from approximately 5,381,001 – 5,474,000 bp on the main bacterial chromosome (CP011972.2; Fig. 3). This deletion site is flanked by the conjugal transfer protein *traG* on both sides, suggesting that this deletion may be mediated by *traG* and, by extension, ICE movement (Fig. 3).

**Figure 3.**
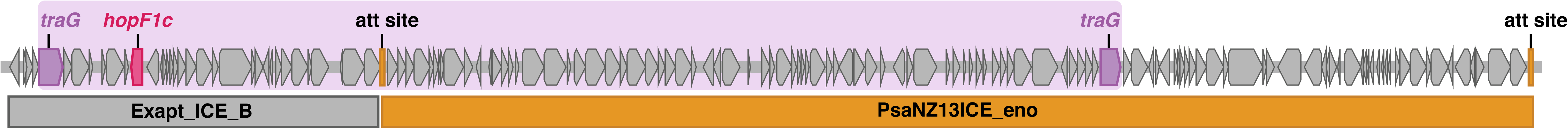
Schematic of the integrative conjugative element PsaNZ13ICE_eno and the neighbouring Exapt_ICE_B. The 93 kb deletion region (5,381,001 – 5,474,000 bp) is highlighted in purple, flanked by traG genes. The effector hopF1c is indicated in pink.

**Figure 4.**
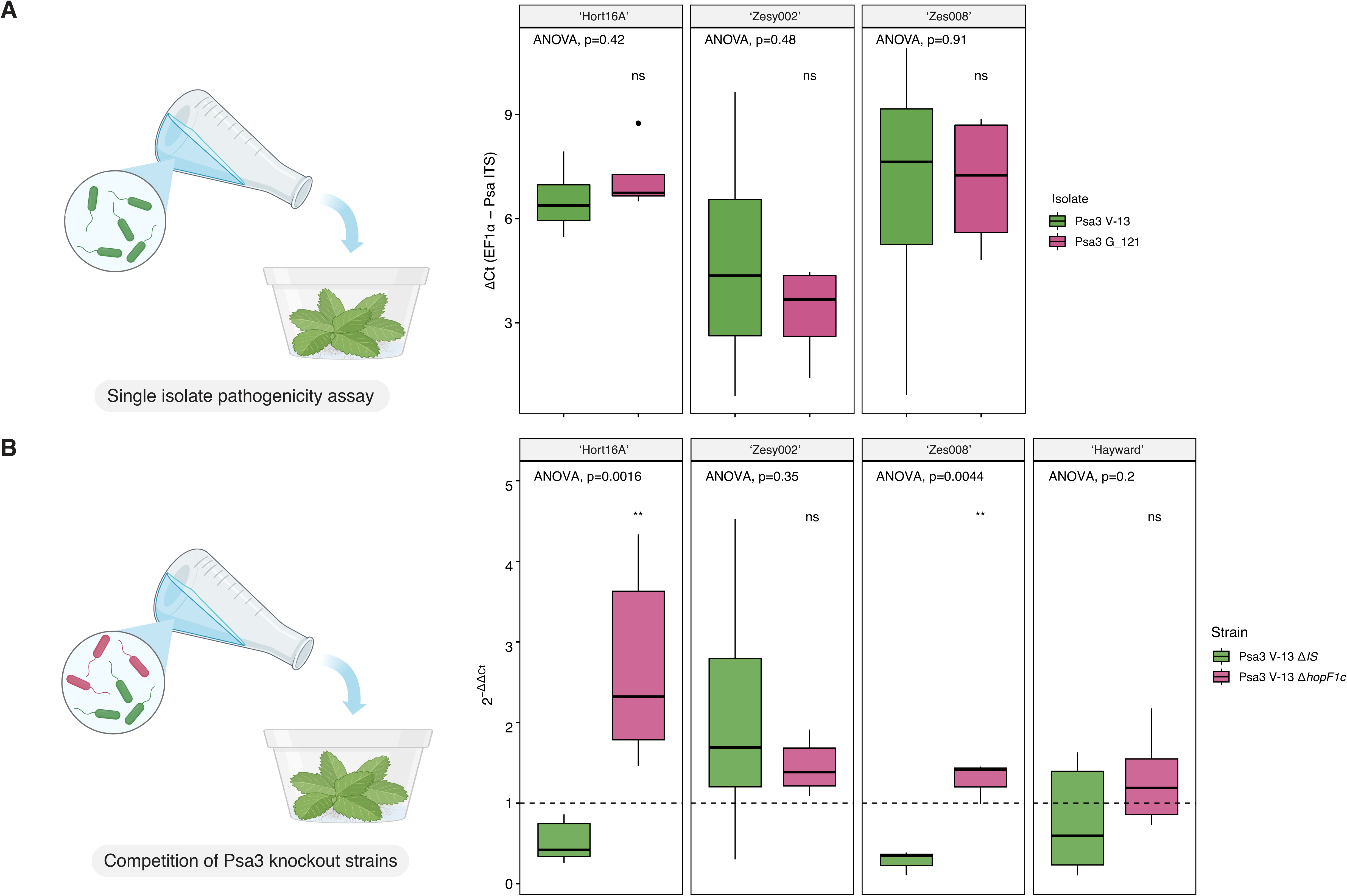
Pathogenicity and competitive fitness of Psa3 hopF1c loss strains on common kiwifruit cultivars. (A) Pathogenicity of Psa3 V-13 and Psa3 V-13 G_121 (lacking hopF1c) on tissue culture Actinidia chinensis var. chinensis cultivars ‘Hort16A’, ‘Zesy002’, and Zes008’. Bacterial pathogenicity was quantified relative to Psa3 V-13 using qPCR ΔCt analysis. Asterisks indicate the statistically significant difference of Welch’s t-test between the indicated strain and wild-type Psa3 V-13, where ns = not significantly different. Horizontal black bars represent the median values. (B). Competitive pathogenicity of ‘wild-type’ Psa3 V-13 ΔIS and Psa3 V-13 ΔhopF1c on tissue culture of four kiwifruit cultivars, Actinidia chinensis var. deliciosa ‘Hayward’ and A. chinensis var. chinensis cultivars ‘Hort16A’, ‘Zesy002’ and ‘Zes008’. Relative quantification was performed at 12 dpi by qPCR ΔΔCt analysis. The boxplots show the relative abundance of effector knockout strains over time, normalised to Psa ITS for each generation and the starting population. Asterisks indicate the statistically significant difference of Welch’s t-test between the indicated strain and Psa3 V-13 ΔIS, where p ≤0.05 (*), p≤0.01 (**) and p ≥0.05 (ns). Horizontal black bars represent the median values.

**Figure 5.**
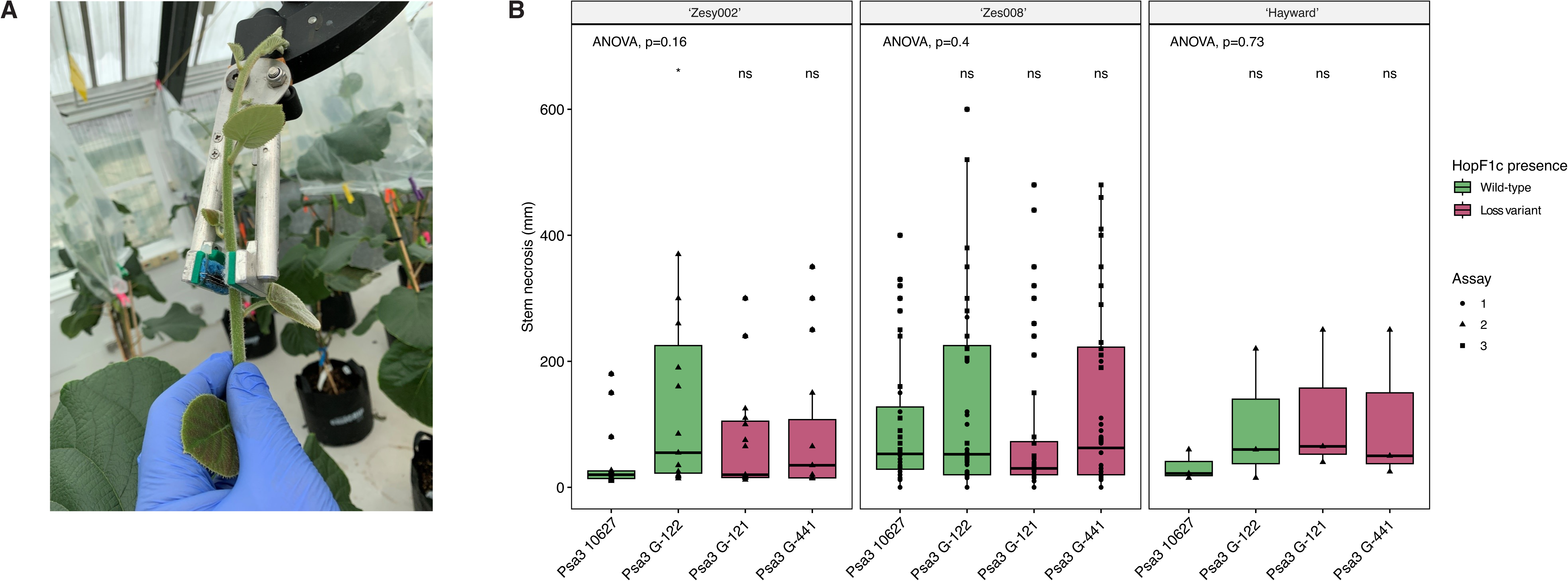
Stem inoculation of Psa3 into potted *Actinidia chinensis* cultivars. (A) Stem inoculation of Psa3 into potted *A. chinensis* plants using a mechanical inoculator. (B) Necrotic lesion length (mm) following stem inoculation of wild-type Psa3 and hopF1c loss isolates into *Actinidia chinensis* var. *chinensis* ‘Zesy002’ and ‘Zes008’ cultivars, and the control *A. chinensis* var. *deliciosa* ‘Hayward’. Asterisks indicate the statistically significant difference of Welch’s t-test between the indicated strain and Psa3 10627, where p ≤0.05 (*) and p ≥0.05 (ns). Horizontal black bars represent the median values.

To assess whether *hopF1c* loss confers a fitness benefit over wild-type Psa3 V-13, the individual pathogenicity of orchard-derived *hopF1c* isolates was tested. Psa3 G_121, an orchard-derived *hopF1c* deletion isolate, showed no significant difference in pathogenicity to wild-type Psa3 V-13 on the *A. chinensis* var. *chinensis* cultivars ‘Hort16A’, ‘Zesy002’, or ‘Zes008’ (Fig. 4A), despite being isolated from ‘Zesy002’. While there was no difference in virulence between Psa3 V-13 and Psa3 G_121 on these *Actinidia* hosts, there was increased growth of both strains on ‘Hort16A’ and ‘Zes008’, compared to ‘Zesy002’ (Fig. 4A). However, when the Psa3 V-13 Δ*hopF1c* knockout strain was tested in a more sensitive competitive fitness assay against a wild-type control (Psa3 V-13 Δ*IS*), Psa3 V-13 Δ*hopF1c* outcompeted wild-type on the *A. chinensis* var. *chinensis* cultivars ‘Hort16A’ and ‘Zes008’, but not ‘Zesy002’ or *Actinidia chinensis* var. *deliciosa* ‘Hayward’ (Fig. 4B). This suggests that, in competition, Psa3 V-13 Δ*hopF1c* may be fitter than wild-type Psa3 on select hosts (Fig. 4B), despite no individual differences in *in planta* growth (Fig. 4A). Pathogenicity assays were repeated on potted plants through stem infection assays (Fig. 5). All inoculated ‘Zes008’ and ‘Zesy002’ shoots developed typical necrotic lesions and there was no significant difference between the *hopF1c* loss and wild-type isolates across all cultivars tested (Fig. 5).

Driven by the discovery and characterisation of these *hopF1c* loss variants – in particular, the competitive fitness of Psa3 V-13 Δ*hopF1c* on select hosts – kiwifruit orchards where *hopF1c* loss isolates had been isolated were resampled. Resampling efforts collected leaves from existing vines, alongside the deployment of young potted trap plants (Fig. S7). Trap plants proved particularly effective for recovering Psa from infected leaf samples, especially when leaves with necrosis were difficult to find in the orchard canopies. Overall, out of a total of 259 Psa isolates collected during this resampling effort, only 6.6% were *hopF1c* loss variants (Table 2). Several *hopF1c* loss isolates were resampled from Orchard B (Table 2). However, none were found at Orchard A, suggesting that the *hopF1c* loss variant has not dominated this site (Table 2). Targeted sampling of other orchards, associated with the original orchards through shared orchard equipment or people movements, also yielded limited numbers of *hopF1c* loss isolates (Table 2). This resampling effort suggests that the spread of *hopF1c* loss isolates is currently limited. In combination with pathogenicity and competitive fitness assays, this suggests that *hopF1c* loss isolates are unlikely to dominate New Zealand’s Psa3 population or cause more severe symptoms than the current population.

**Table 2.** Leaf samples collected from commercial orchards and trap plants during the 2022–23 growing season and the number confirmed with *Pseudomonas syringae* pv. *actinidiae* (Psa). Number and proportion (%) of Psa variants are also shown.

| Source of samples | No. samples taken | No. samples Psa positive | No. of Psa variants identified | Percentage of Psa isolates that were *hopF1c* variants (%) |
| --- | --- | --- | --- | --- |
| Orchard A | 45 | 35 | 0 | 0 |
| Orchard B | 31 | 0 | 0 | 0 |
| Other orchards<sup>a</sup> | 463 | 191 | 11 | 5.8 |
| Trap plants Orchard A | 44 | 27 | 0 | 0 |
| Trap plants Orchard B | 14 | 6 | 6 | 100 |
| Total | 597 | 259 | 17 | 6.56 |

<sup>a</sup>These other orchards were either properties near to Orchards A or B or were associated with these two orchards, in some other way, such as shared orchard equipment or via people movements.

## Discussion

This research sought to characterise genomic changes in New Zealand’s Psa3 population over a 12 year period since its 2010 introduction. Baltrus (2019) suggests that, in *P. syringae* populations, gene gain and loss are likely to be the dominant evolutionary forces, dwarfing the impact of SNPs. Gene gain and loss represent some of the most accessible ways to quickly generate variation in a clonal background, and HGT thus drives the decay of clonality (Gogarten et al., 2002; Puigbò et al., 2014). In keeping with this, the acquisition and loss of mobile genetic elements (MGEs) are significant drivers of variation in New Zealand’s Psa3 population, compared to the negligible impact of *de novo* mutations. The most significant variation in the Psa3 pangenome is due to the excision of the ‘native’ enolase-encoding ICE and the integration of copper-resistance encoding alternatives. Beyond the movement of copper resistance elements, the loss and acquisition of MGEs appear to be limited.

Where genetic element loss has been observed, it has occurred at low frequencies, such as the loss of the urease and achromobactin biosynthetic clusters. Interestingly, the loss of these gene clusters could be driven by either their role in virulence and host adaptation or their redundancy in a high-input orchard environment. Urea is used as a nitrogen source by microbial pathogens of mammals, with urease considered a virulence factor that helps to alkalise acidic environments (Lin et al., 2012; Rutherford, 2014). In diverse microbial pathosystems, urease loss occurs in specialised pathogens to improve host adaptation by reducing toxicity (Chouikha & Hinnebusch, 2014; Baseggio et al., 2022). Alternatively, it could be that kiwifruit orchards with high nutrient inputs select for urease loss, as the potential toxic effect of high ammonium concentrations may negatively impact both Psa3 and its kiwifruit host (Carlini & Polacco, 2008). Urea is also the most common nitrogen fertiliser used in New Zealand (Alizadeh et al., 2017) and is applied in kiwifruit orchards as a fertiliser and to promote leaf drop and degradation in autumn (18 kg/hectare; (Vajari et al., 2018; Max, 2021; Zespri International Ltd, 2022). Achromobactin is an iron-scavenging siderophore (Berti & Thomas, 2009; Owen & Ackerley, 2011), demonstrated to contribute to virulence in *Erwinia chrysanthemi* (Franza et al., 2005) and to epiphytic fitness of *P. syringae* pv. *syringae* (Wensing et al., 2010). However, in some *P. syringae* strains, achromobactin does not contribute to virulence – in these instances, it may be that iron released during infection and plant cell damage renders siderophores obsolete (Owen & Ackerley, 2011). This may explain the loss of these genes in some Psa3 isolates. Alternatively, the iron-scavenging siderophore achromobactin may be rendered redundant through the use of iron-rich supplements in the orchard environment (Berti & Thomas, 2009; Owen & Ackerley, 2011). Finally, a similar situation could explain the loss of an element carrying chemotaxis genes. The high densities of Psa within successfully established lesions on susceptible monocultures may select for the loss of metabolically expensive chemotaxis and motility genes. Alternatively, the addition of surfactant adjuvants to chemical orchard sprays may create an environment where chemotactic motility is less required. Despite their current infrequency, these gene deletions are emerging across independent lineages, suggesting that high-input orchard environments may be one of the few features that Psa is not already well adapted to. Diverse Psa3 isolates have been isolated from kiwifruit orchards in China, despite the fact that Psa could not be found on wild kiwifruit vines there (McCann et al., 2017). If Psa3 originally emerged in Chinese kiwifruit orchards, it could be that these orchards are not as intensively managed as orchards in New Zealand, and the high-input nature of our orchards has provided new selective pressures.

Perhaps the most interesting change in New Zealand’s Psa3 population is the emergence of host-specific effector loss. As observed more generally, the evasion of host recognition has occurred primarily through mobile element excision rather than *de novo* mutagenesis. Limited loss of *hopF1c* was observed in isolates from commercial orchards, in contrast to the more frequent recovery of effector loss isolates from resistant *Actinidia* species in the germplasm collection (Hemara et al., 2022, 2024). The dearth of effector loss in commercial orchards is not necessarily surprising, given that the *A. chinensis* cultivars grown in monoculture are not considered resistant to Psa3, which in turn is unlikely to provide strong selection for effector loss. This raises a further question about what drives *hopF1c* loss in commercial *A. chinensis* orchards. Does *hopF1c* recognition drive *hopF1c* loss (Hemara et al., 2022), even though Psa3 ultimately successfully suppresses HopF1c-triggered immunity in *A. chinensis*? Or could it be that *hopF1c* is simply incidentally excised in the background of frequent Cu^R^ ICE acquisition? To add weight to the former argument, mutation of Psa3 V-13’s ShcF chaperone has reduced HopF1c delivery; this mutation, too, may have been driven by HopF1c recognition as Psa3 circulated in China prior to global spread (Templeton et al., 2015; Jayaraman et al., 2023).

Alternatively, this large deletion may be driven by the combined adaptive fitness of Cu^R^ acquisition and effector loss. Under this scenario, effector loss may be faster than if *hopF1c* was located elsewhere in the genome. Genome context has already been identified as an important factor in Psa effector loss, with the repeated loss of multiple effectors on germplasm driven by repetitive elements around a complex effector locus (Hemara et al., 2022).

Psa3 V-13 Δ*hopF1c* was not fitter than ‘wild-type’ Psa3 V-13 in competition on the dominant commercial cultivars ‘Zesy002’ and ‘Hayward’, despite competitive assays suggesting there might be more subtle differences at play. The role of the *Tn6212* element encoded on the frequently replaced ‘wild-type’ enolase ICE is also of interest when considering drivers of gene loss. *Tn6212* modulates gene expression and may help bacteria adapt to preferred carbon sources across both plant hosts and reservoir environments across the water cycle (Morris et al., 2007, 2008; Colombi et al., 2024). Given their distribution across the star-like phylogeny of Psa strains, the current prevalence of these *hopF1c* effector loss isolates certainly does not constitute a selective sweep of the New Zealand Psa3 population. Escaping the burden of *hopF1c* recognition while simultaneously gaining copper resistance may be beneficial for fitness; however, the benefit of escape from HopF1c recognition may not be sufficient to outcompete other copper resistant isolates in the orchard environment. It remains to be seen whether these effector loss isolates will significantly increase in frequency in current on-orchard Psa3 populations, given more time, or given the introduction of additional kiwifruit genotypes into production.

Comprehensive genome biosurveillance of New Zealand’s once clonal Psa3 population has presented insights into how pathogen populations adapt over time and underscores the importance of carefully considering the deployment of disease management strategies. The emergence of effector loss on both susceptible and resistant kiwifruit vines highlights the importance of managing recurrent pathogen infections, as repeated exposure to host resistance has been repeatedly demonstrated to drive effector loss (Pitman et al., 2005; Arnold et al., 2007; Trivedi & Wang, 2014; Hemara et al., 2022). Whether in monocultures, in the diverse breeding material of germplasm collections, or in wild vines which have escaped the orchard environment, these exposure events provide an opportunity for the pathogen population to break down resistance. The effector loss observed to date also has implications for disease resistance breeding programmes, which seek to introduce resistance genes from Psa-resistant species like *A. arguta* and *A. melanandra* into commercial cultivars. We have demonstrated the recognition of HopF1c, HopAW1a, AvrRpm1a, and HopZ5a across different *Actinidia* species (Hemara et al., 2022, 2024). Should resistant kiwifruit cultivars be deployed as monocultures, the emergence of effector loss isolates at a higher frequency than the current loss of *hopF1c* on susceptible kiwifruit may occur. Further still, as potentially in the case of *hopF1c*, the compounding impact of multiple drivers of resistance breakdown could act in concert to quickly overcome recognition.

A global overreliance on chemical controls, both in horticultural settings and more widely in agricultural and clinical settings, has seen the widespread emergence and spread of factors like Cu resistance genes and Cu resistance-conferring mutations (Meek et al., 2015; Miller et al., 2022; Cella et al., 2023). This also holds true for control technologies used in growing systems, whether that be developing copper resistance (Colombi et al., 2017) or evolving resistance to bacteriophage used as biological control agents (Warring et al., 2022). The loss of biosynthetic gene clusters like achromobactin and urease further demonstrates the potential sensitivity of these populations to chemical inputs. These molecular arms races that drive bacterial adaptation will continue to come to the forefront of effective pathogen control in the coming years. If microbial communities rapidly overcome control strategies upon exposure in the field, the tools we use to manage disease may begin to lose efficacy. Therefore, genome biosurveillance is an increasingly critical tool to detect and quantify adaptations in orchard-based microbial communities. As we seek to diversify horticultural practices to reduce our dependence on chemical controls and move towards more sustainable practices, we must ensure that we monitor and assess the risk of pathogen adaptation to both chemical and biological controls, as well as resistance gene deployment in the field.

## Experimental Procedures

### Kiwifruit vine sampling

Sampling efforts were purposive, based on Psa symptoms including leaf spots and bacterial ooze, as previously described in Hemara et al. (2022, 2024). Symptomatic leaves, buds, shoots and cane were sampled using secateurs sterilised with repeated rinses in 80% ethanol. All plant material was secured in three layers of packaging and stored at approximately 4°C overnight, before transportation back to the laboratory for isolation.

### Commercial orchard sampling & Psa isolation

Sampling was conducted across the Auckland, Waikato, Bay of Plenty, Hawke’s Bay and Northland regions of New Zealand, representative of several of New Zealand’s main kiwifruit growing regions (Table 3). Samples were collected over the spring and summer of each growing season, between November and January. Psa3 isolates, from both this and earlier surveys, from *A. chinensis* var. *chinensis* ‘Zesy002’ and *A. chinensis* var. *deliciosa* ‘Hayward’ vines, as well as the rootstock cultivar *A. chinensis* var. *deliciosa* ‘Bruno’ (Supplementary Table 1). Psa was isolated as described in Hemara et al. (2022). Quantitative PCR (qPCR) was carried out on an Illumina Eco Real-Time PCR platform (Illumina, Melbourne, Australia), following the protocol outlined by Andersen et al. (2017). Single colonies were tested with Psa-ITS F1/R2 PCR primers and primers specific to hopZ5 to identify Psa3 strains.

**Table 3.** Psa3 isolates sampled from commercial kiwifruit orchards across New Zealand. Kiwifruit Growing Region, as classified by Kiwifruit Vine Health, is also indicted.

| Region | Kiwifruit Growing Region | Year |
| --- | --- | --- |
| Auckland | Franklin | 2017, 2018, 2020 |
| Waikato | Waikato | 2017, 2018, 2020, 2022 |
| Bay of Plenty | Tauranga | 2017, 2018, 2020, 2022 |
| Bay of Plenty | Te Puke | 2017, 2018, 2020, 2022 |
| Bay of Plenty | Whakatane | 2017, 2018, 2020, 2022 |
| Hawke's Bay | Hawke's Bay | 2020 |
| Northland | Northland | 2022 |

### DNA extraction and sequencing

DNA was extracted from Psa isolates collected in 2017, 2018, 2019 and 2020, as described in Hemara et al. (2022). For samples collected in 2022, DNA was purified using the Wizard® Genomic DNA Purification Kit (Promega, Wisconsin, United States). Libraries were constructed and sequenced as described in Table 4. Long read sequencing was performed by Auckland Genomics (The University of Auckland, New Zealand) on an Oxford Nanopore Technology (ONT) MINion platform with a R9.4.1 flow cell for select isolates.

**Table 4.**
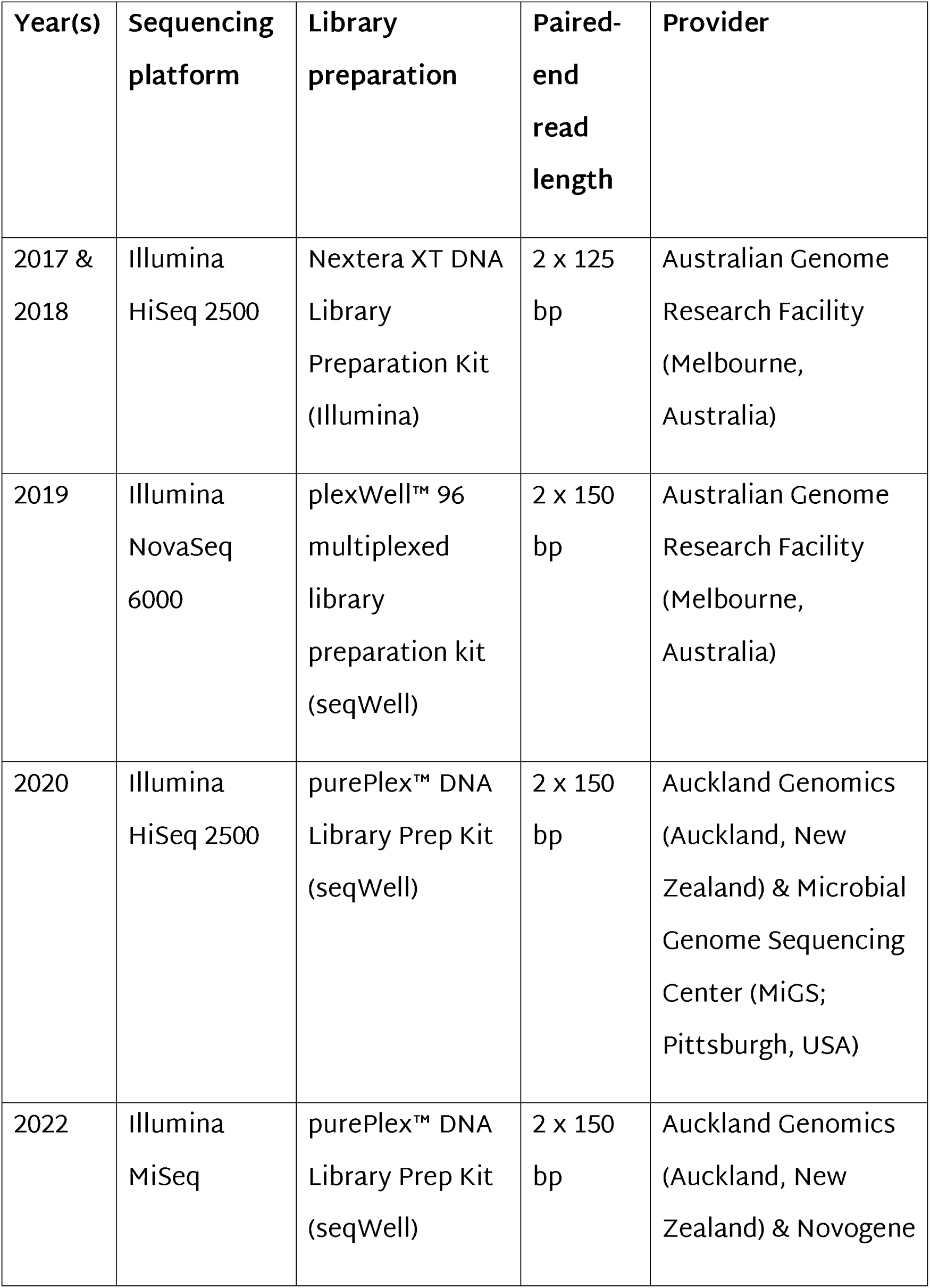
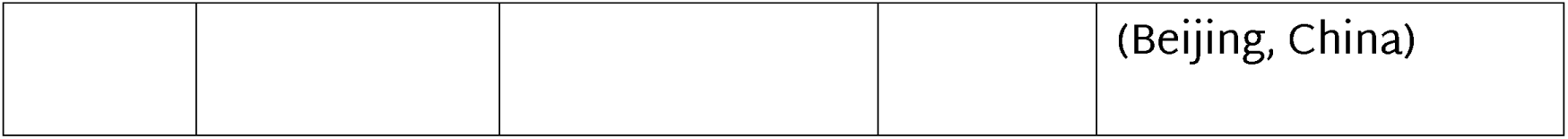
Short-read sequencing platforms used for each isolation year.

### Microbiological methods

All Psa strains were streaked from glycerol stocks onto LB agar supplemented with 12.5 μg/mL nitrofurantoin (Sigma Aldrich, Australia) and 40 μg/mL cephalexin (Sigma Aldrich) for Psa selection. Plates were sealed with parafilm and grown for 48 h at 22°C. LB cultures were grown overnight on a digital orbital shaker at 100 rpm and 22°C.

### Genetically modified bacterial strains

Two genetically modified knockout strains were used in this study: Psa3 V-13 Δ*hopF1c* (Hemara et al., 2022), and Psa3 V-13 Δ*IS*. Psa V-13 Δ*IS* is a derivative of Psa V-13 in which the redundant insertion sequence IS285 was knocked out using methodology described in Jayaraman et al. (2020). Cloning primers are listed in Table 5.

**Table 5.**
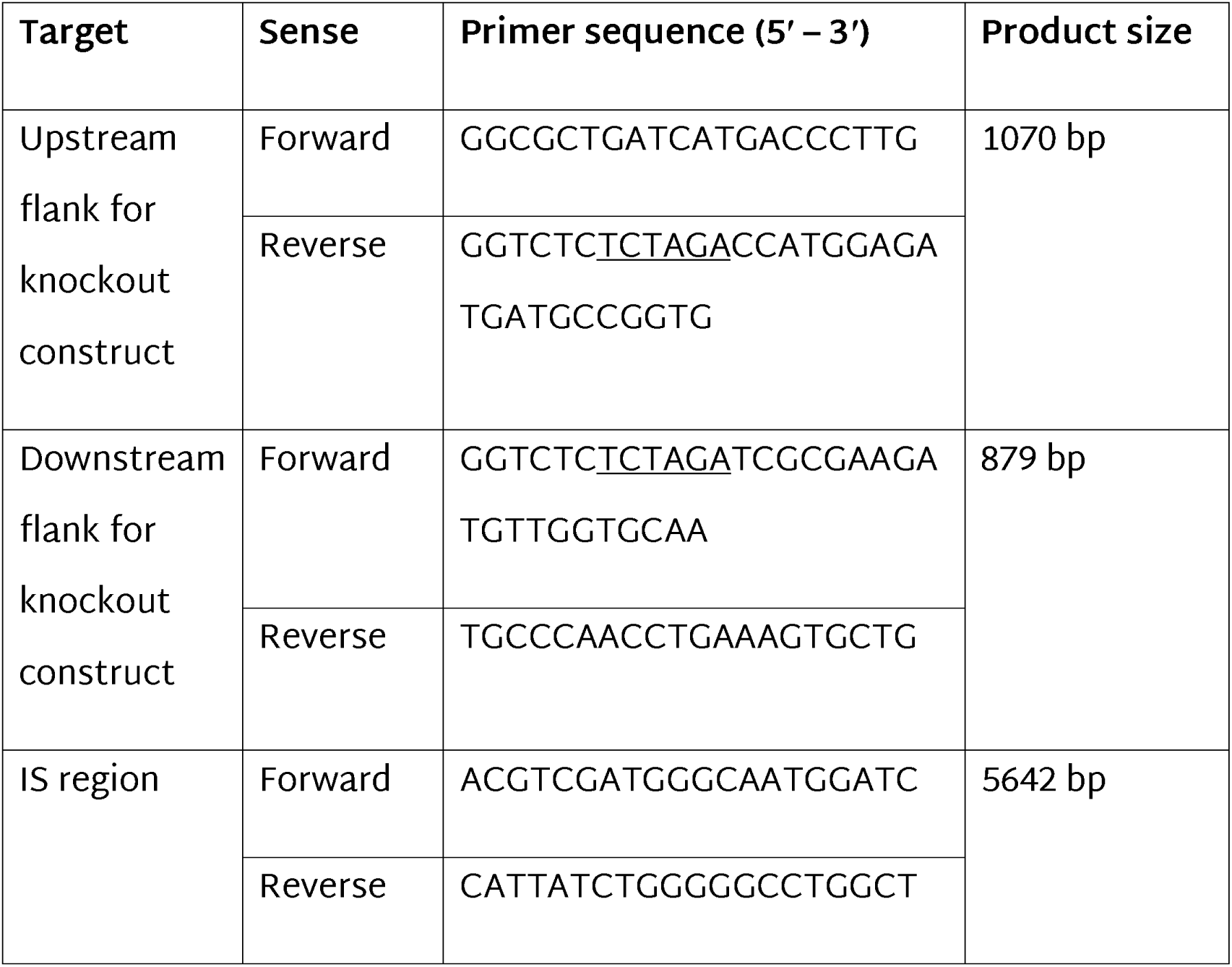
Knockout cloning and confirmation primers used to develop the Psa3 V-13 Δ*IS* knockout strain. The *XbaI* site is underlined.

### Copper resistance testing

Copper resistance was determined by assessing the minimum inhibitory concentration (MIC) of CuSO_4_ required to inhibit bacterial growth, following the protocol described in Colombi et al. (2017).

### In planta pathogenicity and competition assays

#### Tissue culture plantlets

*Actinidia* spp. tissue culture plantlets were supplied by Multiflora Laboratories (Auckland, New Zealand), with the exception of *A. chinensis* var. *chinensis* ‘Zes008’, which was supplied by The New Zealand Institute for Plant and Food Research Ltd (Palmerston North, New Zealand). Each pottle contained three plantlets for *A. chinensis* var. *chinensis* ‘Hort16A’ and ‘Zes008’, and five plantlets for *A. chinensis* var. *chinensis* ‘Zesy002’ and *A. chinensis* var. *deliciosa* ‘Hayward’. *Actinidia* plantlets were grown in 400 mL lidded plastic pottles on half-strength Murashige and Skoog (MS) agar.

#### Flood inoculation

Tissue culture plantlets were flood inoculated with Psa at an OD_600_ of 0.005, using the pathogenicity assay established by McAtee et al. (2018). For single isolate experiments, bacterial growth was quantified at 12 days post-inoculation (dpi) by plate count (Hemara et al., 2022).

#### Competitive *in planta* pathogenicity assays

Psa3 V-13 knockout strains were grown overnight in 5 mL LB and pooled together in equal proportion, with each strain normalised to give an OD_600_ of 0.005 in 500 mL 10 mM MgSO_4_. For the 1:1 effector knockout strain competition assay, Psa V-13 Δ*IS* was used as a control strain and Psa V-13 Δ*hopF1c* represented the *hopF1c* deletion (Hemara et al., 2022). Tissue culture plantlets were flood inoculated, with three replicate pottles per experiment.

Leaf discs were harvested 12 dpi. A 0.8 cm diameter cork borer was used to punch sixteen leaf discs per pottle. Leaf discs were briefly washed in 40 mL of sterile MilliQ H_2_O. Four technical replicates of four leaf discs each were ground in 350 µL sterile 10 mM MgSO_4_ with three 3.5-mm stainless steel beads in a Storm24 Bullet Blender (Next Advance). Samples were ground twice at maximum speed for 1 min. A further 350 µL sterile 10 mM MgSO_4_ was added, and samples were ground at maximum speed for 1 min.

To recover the bacterial population, the resulting leaf homogenate was used to inoculate 50 mL LB supplemented with 12.5 µg/mL nitrofurantoin in a 500 mL conical flask. Leaf homogenate (200 µL) was used for *A. chinensis* var. *chinensis* cultivars, including ‘Hort16A’, ‘Zesy002’ and ‘Zes008’. For *A. chinensis* var. *deliciosa* ‘Hayward’, 300 µL was used. Flasks were shaken on a digital orbital shaker at 100 rpm for 48 h. Aliquots (1 mL) of bacterial culture were sampled after shaking for DNA extraction and long-term glycerol stock storage. DNA was extracted using a Qiagen DNeasy Blood & Tissue Kit (Qiagen, Hilden, Germany), following the Gram-negative bacteria protocol. qPCR was conducted using strain-specific primers to quantify the relative bacterial growth of each strain *in planta*.

#### Potted plant pathogenicity assays

Potted *A. chinensis* var. *chinensis* ‘Zesy002’ and ‘Zes008’ plants were inoculated with four strains of Psa. Suspensions of each Psa strain were delivered directly into the stem using a hand-held mechanical inoculator with two rows of fine stainless-steel needle tips (4 mm long) at a spacing of 1.5 mm (Tahir et al., 2019). This was dipped into the Psa suspension before gently squeezing the needles into a soft and flexible stem section. Stems of large ‘Zes008’ shoots (Assay 1) were inoculated with 2 x 10^8^ CFUs/mL of Psa inoculum approximately 250–400 mm below the growing tip, while other stems (Assays 2 and 3) were inoculated 100–200 mm below the growing tip (due to use of smaller plants). Necrotic stem lesions were measured after 20–26 days and analysed for differences between individual strains and between the wild-type and Psa variant strains.

Two *hopF1c* deletion isolates (Psa3 G_121 and G_441) and two control isolates (Psa3 10627 and G_122) were used. Inoculum was prepared by resuspending strains grown for 2 days on King’s B (KB) medium (King et al., 1954) in sterile water to a final concentration of 1–5 x 10^8^ CFUs/mL. Bacterial concentration was confirmed by plating 10-μL droplets of 1/10 dilutions of the inoculum onto KB plates. The first stem inoculation assay (Assay 1) was carried out using regrowth shoots. For Assays 2 and 3, ‘Zesy002’ and ‘Zes008’ tissue culture plantlets were ex-flasked and potted into 1-L pots, with potted plants 250–500 mm in height at the time of inoculation.

#### Confirmation of Psa from symptomatic tissues

Bacteria were isolated from 12–24 stem samples for each of the Psa strains used in each of the plant assays by macerating necrotic stems and their surrounding tissue in 300 µL of sterile distilled water. Fifty–100 µL of the macerate was streaked on agar plates of KB and KB medium supplemented with 1.5% boric acid and cephalexin 80 mg/L (KBC); both media were also supplemented with 1% cycloheximide. The KB plates were incubated at 28°C for 48 h and the KBC plates were incubated for 7 d before purifying colonies showing a morphology similar to that of Psa. The identity of the bacteria was confirmed by duplex PCR using the primer pair Psa F1/R2 (Rees-George et al., 2010) and the primers Psa-*hopF1c*-F and Psa-*hopF1c*-R.

#### Spread of new Psa variant in orchards and use of trap plants

During the spring of 2022, Kiwifruit Vine Health (KVH) led delimiting surveillance studies to manage the movement of risk goods and reduce risk to neighbouring properties. Leaves with necrotic spotting from Orchards A and B where Psa variants had been discovered, and from nearby and associated orchards (e.g. through shared orchard equipment), were sent to Hill Laboratories (Hamilton, New Zealand) to recover Psa and determine if the Psa variant was present.

Potted kiwifruit trap plants of *A. chinensis* var. *deliciosa* ‘Hayward’ or ‘Bruno’ were placed in Orchards A and B to recover Psa, as it was difficult finding leaves with Psa leaf necrosis. Twenty trap plants were placed in Orchard A on two occasions (27 September to 19 October 2022) and exposed for 7 and 13 d, respectively, before being returned to a covered outdoor area for 2–3 weeks to allow symptom development. In Orchard B, trap plants were set up on 21 November 2022 and leaf sampling was carried out *in situ* after 21 d & sent to Hill Laboratories for processing.

#### DNA extraction

The Qiagen DNAeasy Blood & Tissue kit was used for DNA extractions from LB culture, following the protocol for gram-negative bacteria (Qiagen, Hilden, Germany). DNA samples were diluted 1/10 before being used as templates for quantitative PCR.

#### Quantitative PCR

Real-time qPCR was performed using an Illumina Eco Real-Time PCR platform, following the protocol developed by Andersen et al. (2017). Bacterial growth was assessed by relative quantification, for the wildtype-like strain (ΔIS) using forward (5ʹ-ACTACTTCACCCAGGACCTG-3ʹ) and reverse (5ʹ-CGTTTGCACCAACATCTTCG-3ʹ) primers, and for the Δ*hopF1c* knockout strain using forward (5ʹ-TCCACAGCATGACCAACA-GT-3ʹ) and reverse (5ʹ-TGCGGTCGATCAAAATCTCTAGA-3ʹ) primers. The cycle threshold (Ct) value for each knockout primer pair was normalised, using the ΔΔCt method, to the Psa ITS Ct value, and then to the ΔCt values for the original inoculum/pool. Relative quantification values were visualised as 2^-ΔΔCt^, with each knockout strain normalised to Psa ITS for each generation and the starting population.

For the identification of *hopF1c* isolates, primers that flanked the *hopF1c* gene were designed using Geneious R11 software (https://www.geneious.com; Biomatters). Forward (5ʹ-TGTGGTA-CTTCTGGCTCTCATCA-3ʹ) and reverse (5ʹ-TCGTCCACTACCTGCGCT-3ʹ) primers amplified a 602 bp fragment in wild-type strains of Psa but failed to amplify a band from*hopF1c* loss variants.

### Bioinformatic methods

#### Psa3 genome assembly and pangenome analysis

Quality control reports for the raw sequencing reads were generated using FastQC (version 0.11.7; Andrews, 2010). Paired-end reads were assembled using SPAdes (version 3.14.0; Bankevich et al., 2012) and shovill (version 0.9.0; https://github.com/tseemann/shovill).

Trycycler (version 0.5.4) was used to curate consensus long-read assemblies (Wick et al., 2021), using the Flye (version 2.9.2; Kolmogorov et al., 2019) and Canu (version 2.2; Koren et al., 2017) assemblers. Unicycler (version 0.5.0) was used to generate hybrid assemblies (Wick et al., 2017).

Contigs were annotated with Prokka (version 1.3; Seemann, 2014) using the Psa3 V-13 (ICMP 18884) protein model. PADLOC was used to identify antiphage defence systems (version 2.0.0; Payne et al., 2022). Roary (version 3.13.0; Page et al., 2015) and Panaroo (version 1.3.0; Tonkin-Hill et al., 2020) were used for pangenome analysis. Phandango (Hadfield et al., 2018) and pagoo (version 0.3.17; Ferrés & Iraola, 2021) were used for pangenome visualisation. Mandrake (version 1.2.2; Lees et al., 2022) was used to produce stochastic cluster embedding visualisations from pangenome gene presence/absence data.

#### Psa3 variant calling

Snippy (version 4.6.0) was used to map Psa3 reads to the Psa3 V-13 reference genome and snippy-core was used to produce a core SNP alignment (Seemann, 2015). New Zealand Psa3 isolates from both commercial kiwifruit orchards (Supplementary Table 1) and germplasm collections (Hemara et al., 2022; 2024) were used. Gubbins (version 2.4.1) identified recombinant regions in this alignment, producing a filtered alignment (Croucher et al., 2015). RAxML (version 8.2.12; –f a –# 100 –m GTRCAT) was used to generate a maximum-likelihood phylogenetic tree with 100 bootstrap replicates (Stamatakis, 2014). The phylogeny and associated metadata were visualised with the R package ggtree (version 2.2.4; Yu et al., 2017). Only bootstrap support values of 50 or above were visualised.

Unmapped reads, as output by Snippy, were assembled using SPAdes (version 3.14.0; Bankevich et al., 2012) and annotated with Prokka (version 1.3; Seemann, 2014). Magic-BLAST (Boratyn et al., 2019) was used to build custom BLAST databases of known effectors as defined by Dillon et al. (2019), ICEs (Colombi et al., 2017; Poulter et al., 2018), and plasmids to BLAST the assembled unmapped contigs with BLAST+ (version 2.10.1+; Camacho et al., 2009). Novel elements were identified using the NCBI Nucleotide BLAST web interface (Johnson et al., 2008). Gene deletions were identified from bam format read alignments by CNVnator (version 0.4.1; Abyzov et al., 2011) and compared to pangenome gene presence/absence data.

#### Data visualisation & statistical analysis

Statistical analysis was conducted in R (R Core Team, 2024), and figures were produced using the packages ggplot2 (Wickham, 2016) and ggpubr (Kassambara, 2017). Plots were exported from R as PDF files and prepared for publication in Adobe Illustrator (Adobe Inc.). Post-hoc statistical tests were conducted using the ggpubr (version 0.3.0) and agricolae (version 1.3) packages (de Mendiburu, 2017; Kassambara, 2017). The stats_compare_means() function from the ggpubr package was used to calculate omnibus one-way analysis of variance (ANOVA) statistics to identify statistically significant differences across all treatment groups (Kassambara, 2017). For normally distributed populations, Welch’s t-test was used to conduct pair-wise parametric t-tests between an indicated group and a designated reference (Kassambara, 2017). The HSD.test() function from the agricolae package was used to calculate Tukey’s Honest Significant Difference (de Mendiburu, 2017).

Graphical schematics were made in BioRender (https://www.biorender.com/).

## Acknowledgements

The authors gratefully thank: Dr Honour McCann for bioinformatics guidance; Dr Shahjahan Kabir and Teiarere Stephens for trap plant assistance; and Drs Erik Rikkerink, Kirstin Wurms, Philip Elmer, Tony Reglinski and Chandan Pal for reviewing iterations of this manuscript. The authors wish to acknowledge funding provided by Zespri Group Ltd and the collaborative efforts of Kiwifruit Vine Health, Zespri, Plant & Food Research, orchard managers, and kiwifruit growers. LMH would like to thank the University of Auckland for a Doctoral Scholarship. The authors wish to acknowledge the use of New Zealand eScience Infrastructure (NeSI) high performance computing facilities, consulting support and training services as part of this research. New Zealand’s national facilities are provided by NeSI and funded jointly by NeSI’s collaborator institutions and through the Ministry of Business, Innovation & Employment’s Research Infrastructure programme. URL https://www.nesi.org.nz.

## Data availability statement

Sequence data are available in the GenBank Nucleotide Database (https://www.ncbi.nlm.nih.gov/genbank/) and Sequence Read Archive (https://www.ncbi.nlm.nih.gov/sra) under BioProjects PRJNA826130, PRJNA826136, PRJNA1165291, PRJNA826129, PRJNA826132, PRJNA826143, and PRJNA1165295.

## Notes

### Competing Interest Statement

The authors have declared no competing interest.

