## Supporting Information for "Genomic biosurveillance of the kiwifruit pathogen Pseudomonas syringae pv. actinidiae biovar 3 reveals adaptation to selective pressures in New Zealand orchards"

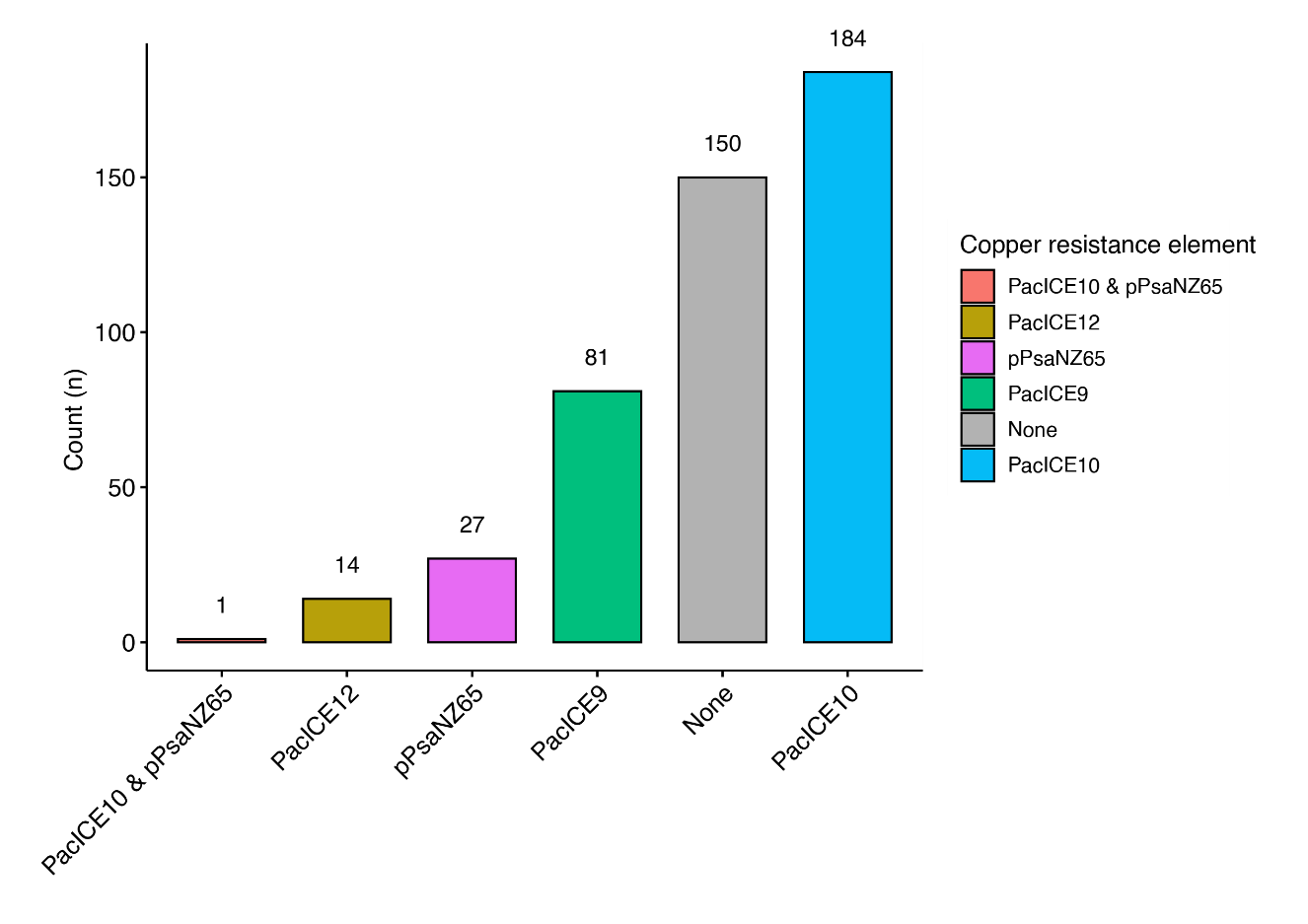

**Supplementary Figure 1. Copper resistance elements found in New Zealand Psa3 isolates.**

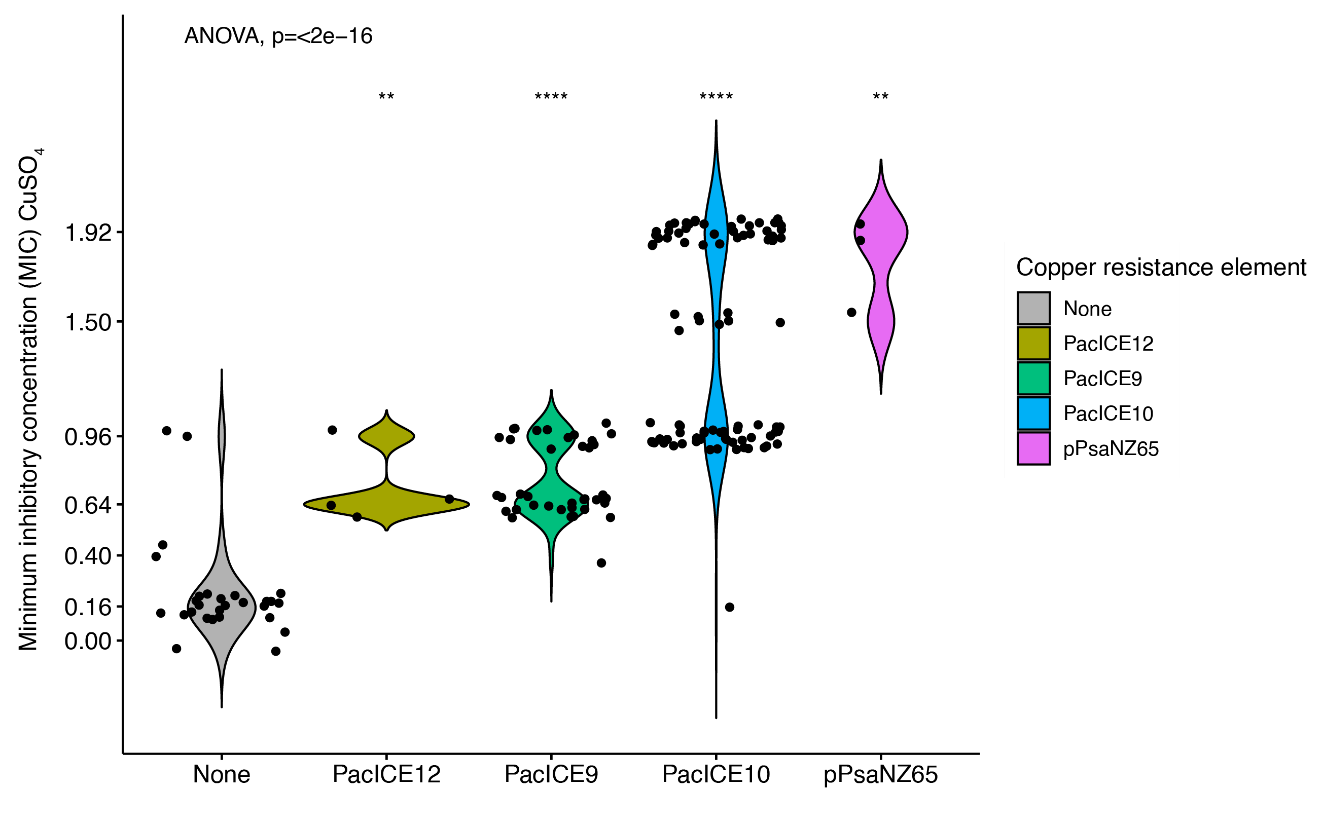

**Supplementary Figure 2. Minimum inhibitory concentration (MIC; mM) of CuSO_4_ by copper resistance element.**

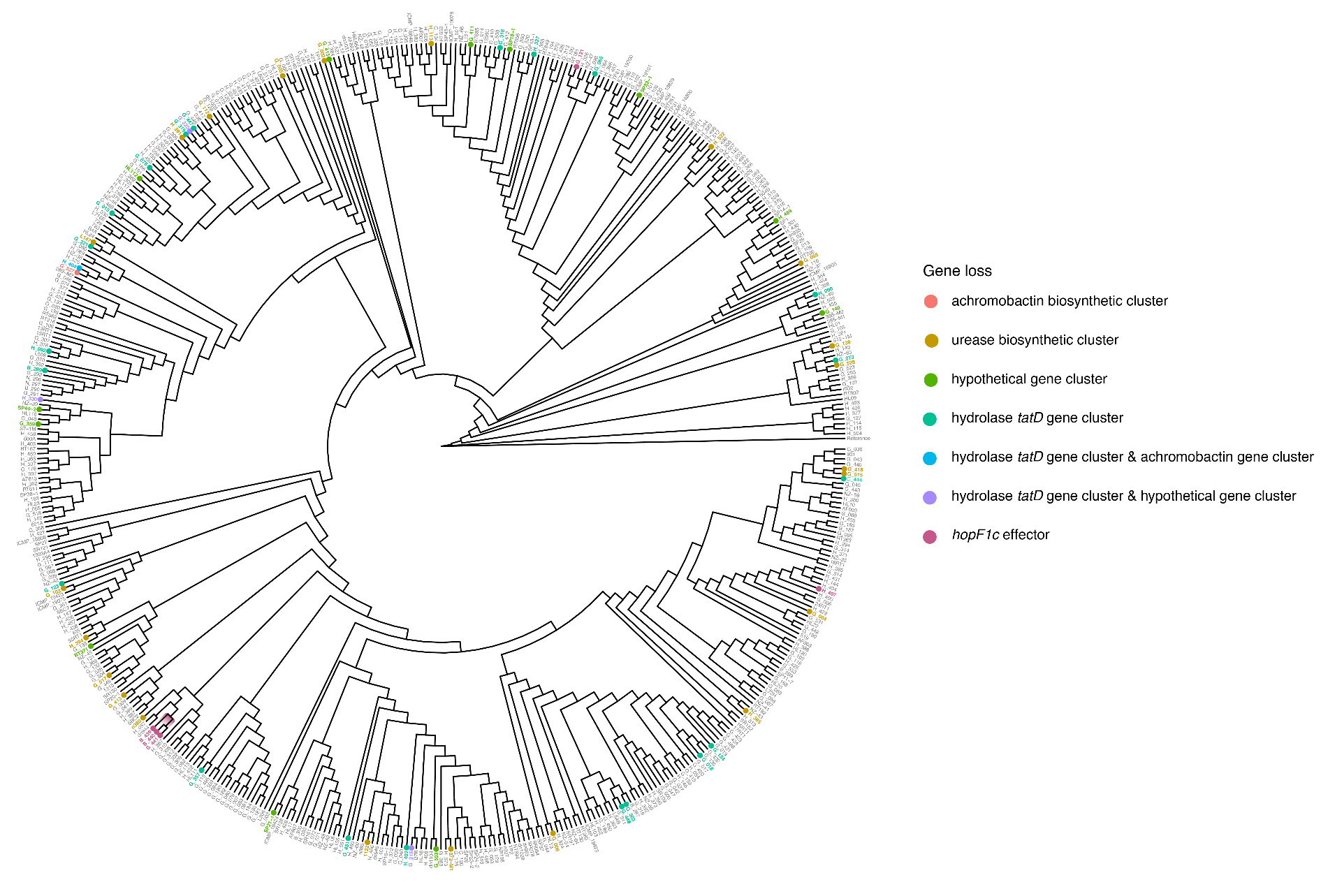
**Supplementary Figure 3. Core SNP cladogram of commercial New Zealand Psa3 isolates with key gene loss events highlighted.** A core SNP phylogeny of New Zealand Psa3 isolates was produced with Snippy (version 4.6.0) relative to the reference Psa3 V-13. Gene loss events are highlighted, as identified by CNVnator (version 0.4.1; Abyzov et al., 2011) and Panaroo (version 1.3.0; Tonkin-Hill et al., 2020).

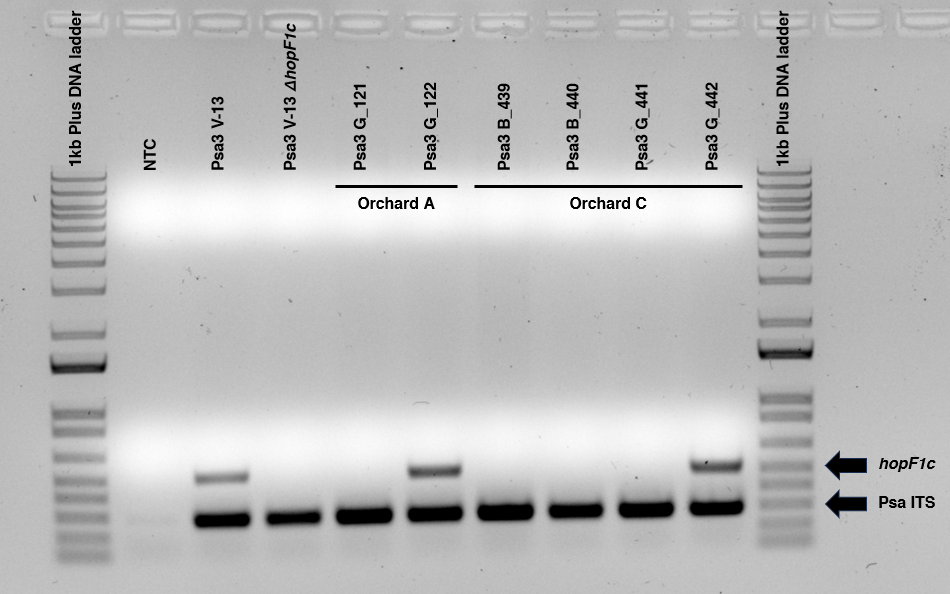

**Supplementary Figure 4: Multiplex PCR confirms hopF1c has been lost from Psa3 G_035, Psa3 B_439, Psa3 B_440, and Psa3 G_441.** Expected hopF1c and Psa ITS band sizes are indicated on the 1 Kb Plus DNA ladder (ThermoFisher Scientific, New Zealand).

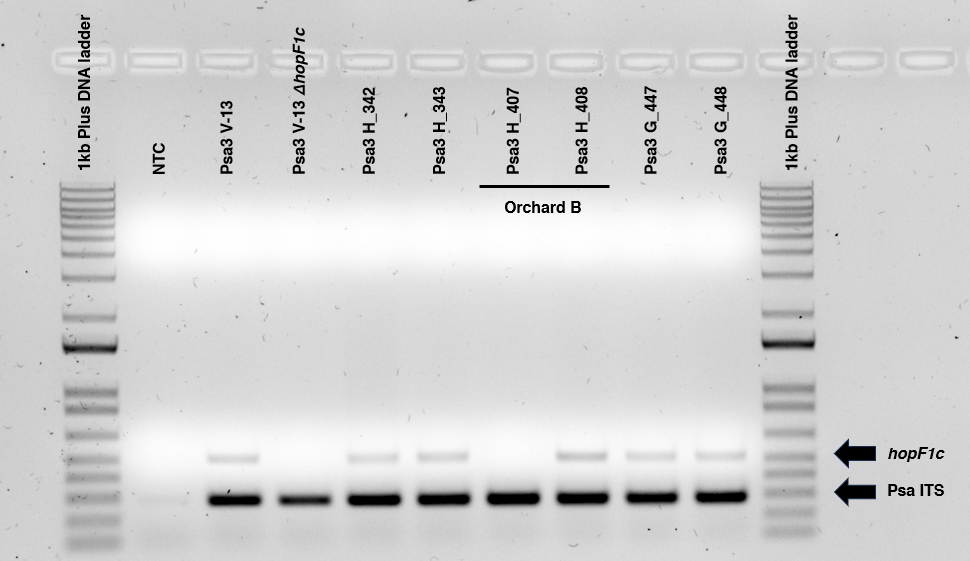

**Supplementary Figure 5: Multiplex PCR confirms hopF1c has been lost from Psa3 H_407.** Expected hopF1c and Psa ITS band sizes are indicated on the 1 Kb Plus DNA ladder (ThermoFisher Scientific, New Zealand).

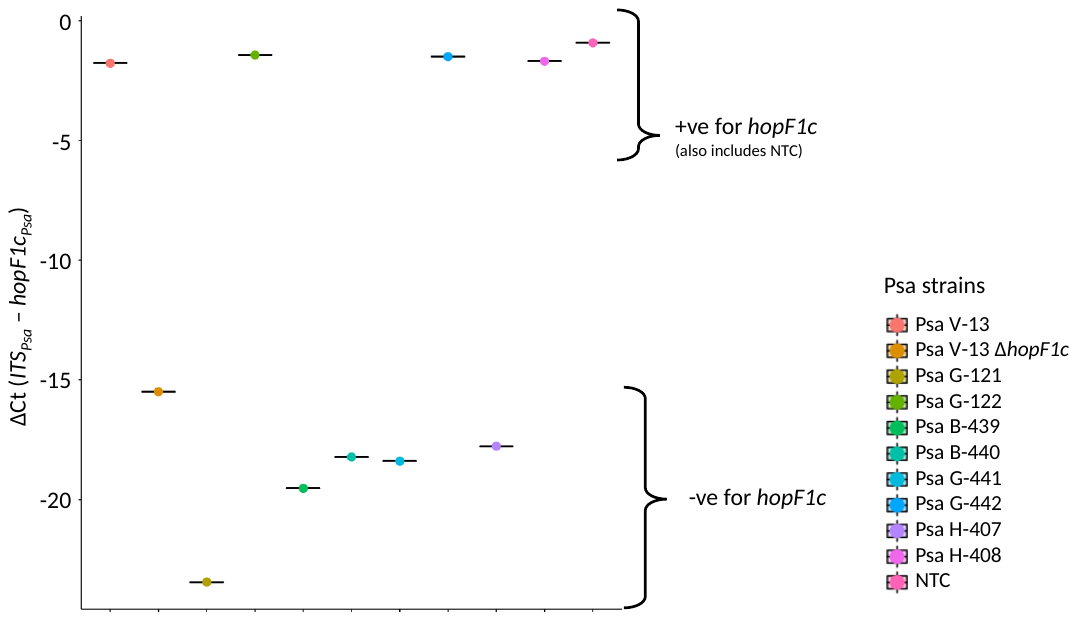

**Supplementary Figure 6. Quantitative PCR confirms hopF1c has been lost from orchard-derived Psa3 isolates.** *hopF1c* presence was assessed using qPCR ΔCt analysis, using the ITS primers from Rees-George et al. (2010) and those designed to amplify *hopF1c*. NTC = no template control.

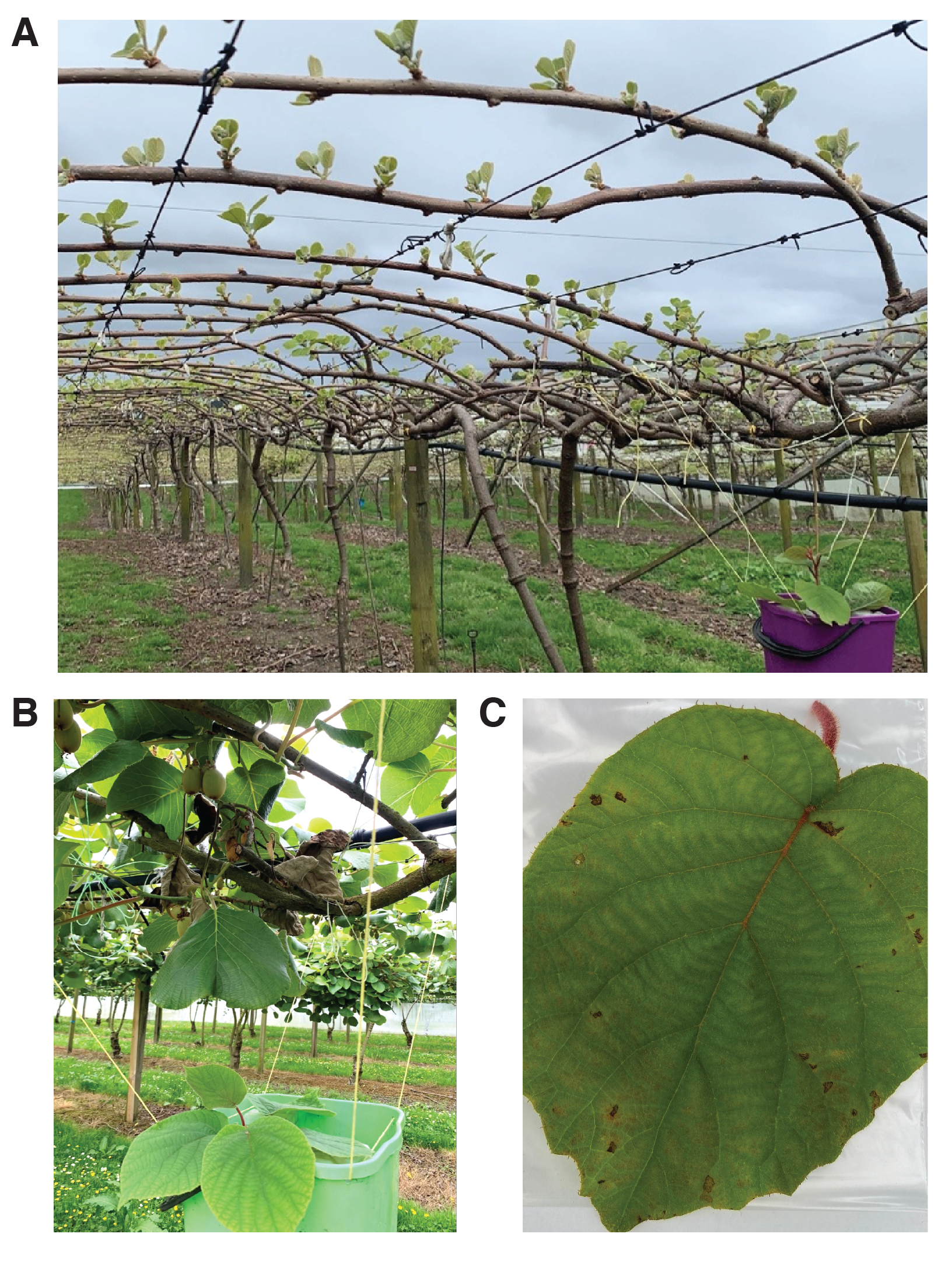

**Supplementary Figure 7. Trap plants deployed in kiwifruit orchards of interest to survey the relative abundance of Psa3 *hopF1c* loss variants.** (A) *Actinidia chinensis* var. *deliciosa* ‘Hayward’ trap plants deployed in Orchard A in spring, with the orchard showing early season growth. (B) *A. chinensis* var. *chinensis* ‘Hayward’ trap plants suspended beneath the orchard canopy in Orchard B, located directly under typical shoot dieback symptoms observed during set-up. (C) Example of a trap plant leaf which has developed typical Psa lesion symptoms after orchard deployment.

**Supplementary Table 1. New Zealand Psa3 isolates from commercial kiwifruit (Actinidia chinensis) orchards used in this study.** For isolates originating from this study, the letter codes denote host of isolation, with G representing A. chinensis var. chinensis ‘Zesy002’, H representing A. chinensis var. deliciosa ‘Hayward’, and B representing A. chinensis var. deliciosa ‘Bruno’.

| Isolate | Year | Bioproject | Source |
| --- | --- | --- | --- |
| Psa3 ICMP 18800 | 2010 | PRJNA527140 | University of Otago |
| Psa3 ICMP 18805 | 2010 | PRJNA527140 | University of Otago |
| Psa3 ICMP 18808 | 2010 | PRJNA527140 | University of Otago |
| Psa3 ICMP 19075 | 2010 | PRJNA527140 | University of Otago |
| Psa3 ICMP 19101 | 2010 | PRJNA527140 | University of Otago |
| Psa3 NZ-35 | 2010 | PRJNA359702 | (McCann et al., 2017) |
| Psa3 NZ-37 | 2010 | PRJNA359702 | (McCann et al., 2017) |
| Psa3 V-13 (ICMP 18884) | 2010 | PRJNA71845 | (McCann et al., 2013; Templeton et al., 2015) |
| Psa3 ICMP 18839 | 2011 | PRJNA527140 | University of Otago |
| Psa3 ICMP 18848 | 2011 | PRJNA527140 | University of Otago |
| Psa3 ICMP 18875 | 2011 | PRJNA527140 | University of Otago |
| Psa3 ICMP 19076 | 2011 | PRJNA527140 | University of Otago |
| Psa3 ICMP 19200 | 2011 | PRJNA527140 | University of Otago |
| Psa3 ICMP 19423 | 2011 | PRJNA527140 | University of Otago |
| Psa3 ICMP 19424 | 2011 | PRJNA527140 | University of Otago |
| Psa3 NZ-49 | 2011 | PRJNA359702 | (McCann et al., 2017) |
| Psa3 TP1 | 2011 | PRJNA527140 | University of Otago |
| Psa3 TP2 | 2011 | PRJNA527140 | University of Otago |
| Psa3 TP22 | 2011 | PRJNA527140 | University of Otago |
| Psa3 TP31 | 2011 | PRJNA527140 | University of Otago |
| Psa3 TP61 | 2011 | PRJNA527140 | University of Otago |
| Psa3 13RT1 | 2012 | PRJNA527140 | University of Otago |
| Psa3 15RT1 | 2012 | PRJNA527140 | University of Otago |
| Psa3 16RT1 | 2012 | PRJNA527140 | University of Otago |
| Psa3 17RT1 | 2012 | PRJNA527140 | University of Otago |
| Psa3 18RT1 | 2012 | PRJNA527140 | University of Otago |
| Psa3 24RT1 | 2012 | PRJNA527140 | University of Otago |
| Psa3 377 | 2012 | PRJNA527140 | University of Otago |
| Psa3 387 | 2012 | PRJNA527140 | University of Otago |
| Psa3 405 | 2012 | PRJNA527140 | University of Otago |
| Psa3 421 | 2012 | PRJNA527140 | University of Otago |
| Psa3 55RT1 | 2012 | PRJNA527140 | University of Otago |
| Psa3 5RT1 | 2012 | PRJNA527140 | University of Otago |
| Psa3 723 | 2012 | PRJNA527140 | University of Otago |
| Psa3 729 | 2012 | PRJNA527140 | University of Otago |
| Psa3 901 | 2012 | PRJNA527140 | University of Otago |
| Psa3 936 | 2012 | PRJNA527140 | University of Otago |
| Psa3 AC507 | 2012 | PRJNA527140 | University of Otago |
| Psa3 NZ-46 | 2012 | PRJNA359702 | (McCann et al., 2017) |
| Psa3 B4A | 2013 | PRJNA527140 | University of Otago |
| Psa3 B7B | 2013 | PRJNA527140 | University of Otago |
| Psa3 L2B | 2013 | PRJNA527140 | University of Otago |
| Psa3 L3AB | 2013 | PRJNA527140 | University of Otago |
| Psa3 NZ-48 | 2013 | PRJNA359702 | (McCann et al., 2017) |
| Psa3 AF807 | 2014 | PRJNA527140 | University of Otago |
| Psa3 AF813 | 2014 | PRJNA527140 | University of Otago |
| Psa3 AF862 | 2014 | PRJNA527140 | University of Otago |
| Psa3 AF868 | 2014 | PRJNA527140 | University of Otago |
| Psa3 AF903 | 2014 | PRJNA527140 | University of Otago |
| Psa3 AF907 | 2014 | PRJNA527140 | University of Otago |
| Psa3 NZ-38 | 2014 | PRJNA359702 | (McCann et al., 2017) |
| Psa3 NZ-39 | 2014 | PRJNA359702 | (McCann et al., 2017) |
| Psa3 NZ-40 | 2014 | PRJNA359702 | (McCann et al., 2017) |
| Psa3 NZ-41 | 2014 | PRJNA359702 | (McCann et al., 2017) |
| Psa3 NZ-42 | 2014 | PRJNA359702 | (McCann et al., 2017) |
| Psa3 NZ-43 | 2014 | PRJNA359702 | (McCann et al., 2017) |
| Psa3 NZ-45 | 2014 | PRJNA359702 | (McCann et al., 2017) |
| Psa3 NZ-47 | 2014 | PRJNA359702 | (McCann et al., 2017) |
| Psa3 NZ-54 | 2014 | PRJNA359702 | (McCann et al., 2017) |
| Psa3 NZ-62 | 2014 | PRJNA359702 | (McCann et al., 2017) |
| Psa3 RT130 | 2014 | PRJNA527140 | University of Otago |
| Psa3 RT167 | 2014 | PRJNA527140 | University of Otago |
| Psa3 RT181 | 2014 | PRJNA527140 | University of Otago |
| Psa3 RT182 | 2014 | PRJNA527140 | University of Otago |
| Psa3 RT216 | 2014 | PRJNA527140 | University of Otago |
| Psa3 RT261 | 2014 | PRJNA527140 | University of Otago |
| Psa3 RT363 | 2014 | PRJNA527140 | University of Otago |
| Psa3 RT371 | 2014 | PRJNA527140 | University of Otago |
| Psa3 RT383 | 2014 | PRJNA527140 | University of Otago |
| Psa3 dn1005 | 2015 | PRJNA527140 | University of Otago |
| Psa3 dn1028 | 2015 | PRJNA527140 | University of Otago |
| Psa3 dn1031 | 2015 | PRJNA527140 | University of Otago |
| Psa3 dn1104 | 2015 | PRJNA527140 | University of Otago |
| Psa3 dn935 | 2015 | PRJNA527140 | University of Otago |
| Psa3 dn944 | 2015 | PRJNA527140 | University of Otago |
| Psa3 dn946 | 2015 | PRJNA527140 | University of Otago |
| Psa3 dn985 | 2015 | PRJNA527140 | University of Otago |
| Psa3 NZ-59 | 2015 | PRJNA359702 | (McCann et al., 2017) |
| Psa3 NZ-60 | 2015 | PRJNA359702 | (McCann et al., 2017) |
| Psa3 NZ-63 | 2015 | PRJNA339324 | (Colombi et al., 2017) |
| Psa3 RT594 | 2015 | PRJNA527140 | University of Otago |
| Psa3 RT652 | 2015 | PRJNA527140 | University of Otago |
| Psa3 RT656 | 2015 | PRJNA527140 | University of Otago |
| Psa3 RT685 | 2015 | PRJNA527140 | University of Otago |
| Psa3 RT786 | 2015 | PRJNA527140 | University of Otago |
| Psa3 RT802 | 2015 | PRJNA527140 | University of Otago |
| Psa3 RT811 | 2015 | PRJNA527140 | University of Otago |
| Psa3 RT812 | 2015 | PRJNA527140 | University of Otago |
| Psa3 RT849 | 2015 | PRJNA527140 | University of Otago |
| Psa3 RT862 | 2015 | PRJNA527140 | University of Otago |
| Psa3 SR078 | 2015 | PRJNA527140 | University of Otago |
| Psa3 SR084 | 2015 | PRJNA527140 | University of Otago |
| Psa3 SR121 | 2015 | PRJNA527140 | University of Otago |
| Psa3 SR130 | 2015 | PRJNA527140 | University of Otago |
| Psa3 SR138 | 2015 | PRJNA527140 | University of Otago |
| Psa3 SR140 | 2015 | PRJNA527140 | University of Otago |
| Psa3 SR152 | 2015 | PRJNA527140 | University of Otago |
| Psa3 SR161 | 2015 | PRJNA527140 | University of Otago |
| Psa3 SR197 | 2015 | PRJNA527140 | University of Otago |
| Psa3 SR198 | 2015 | PRJNA527140 | University of Otago |
| Psa3 SR199 | 2015 | PRJNA527140 | University of Otago |
| Psa3 HL01 | 2016 | PRJNA527140 | University of Otago |
| Psa3 HL09 | 2016 | PRJNA527140 | University of Otago |
| Psa3 HL10 | 2016 | PRJNA527140 | University of Otago |
| Psa3 HL11 | 2016 | PRJNA527140 | University of Otago |
| Psa3 HL13 | 2016 | PRJNA527140 | University of Otago |
| Psa3 HL16 | 2016 | PRJNA527140 | University of Otago |
| Psa3 HL17 | 2016 | PRJNA527140 | University of Otago |
| Psa3 HL18 | 2016 | PRJNA527140 | University of Otago |
| Psa3 HL19 | 2016 | PRJNA527140 | University of Otago |
| Psa3 HL20 | 2016 | PRJNA527140 | University of Otago |
| Psa3 HL22 | 2016 | PRJNA527140 | University of Otago |
| Psa3 HL23 | 2016 | PRJNA527140 | University of Otago |
| Psa3 HL24 | 2016 | PRJNA527140 | University of Otago |
| Psa3 NZ-64 | 2016 | PRJNA339325 | (Colombi et al., 2017) |
| Psa3 NZ-65 | 2016 | PRJNA339326 | (Colombi et al., 2017) |
| Psa3 NZ-66 | 2016 | PRJNA339327 | (Colombi et al., 2017) |
| Psa3 1122 | 2017 | PRJNA527140 | University of Otago |
| Psa3 1215 | 2017 | PRJNA527140 | University of Otago |
| Psa3 1256C | 2017 | PRJNA527140 | University of Otago |
| Psa3 1393A4 | 2017 | PRJNA527140 | University of Otago |
| Psa3 373-M1 | 2017 | PRJNA527140 | University of Otago |
| Psa3 385-M1 | 2017 | PRJNA527140 | University of Otago |
| Psa3 388-M2 | 2017 | PRJNA527140 | University of Otago |
| Psa3 389-M2 | 2017 | PRJNA527140 | University of Otago |
| Psa3 600A | 2017 | PRJNA527140 | University of Otago |
| Psa3 601A | 2017 | PRJNA527140 | University of Otago |
| Psa3 604B | 2017 | PRJNA527140 | University of Otago |
| Psa3 G_001 | 2017 | PRJNA826129 | This study |
| Psa3 G_002 | 2017 | PRJNA826129 | This study |
| Psa3 G_003 | 2017 | PRJNA826129 | This study |
| Psa3 G_004 | 2017 | PRJNA826129 | This study |
| Psa3 G_005 | 2017 | PRJNA826129 | This study |
| Psa3 G_006 | 2017 | PRJNA826129 | This study |
| Psa3 G_007 | 2017 | PRJNA826129 | This study |
| Psa3 G_008 | 2017 | PRJNA826129 | This study |
| Psa3 G_009 | 2017 | PRJNA826129 | This study |
| Psa3 G_012 | 2017 | PRJNA826129 | This study |
| Psa3 G_013 | 2017 | PRJNA826129 | This study |
| Psa3 G_014 | 2017 | PRJNA826129 | This study |
| Psa3 G_015 | 2017 | PRJNA826129 | This study |
| Psa3 G_016 | 2017 | PRJNA826129 | This study |
| Psa3 G_017 | 2017 | PRJNA826129 | This study |
| Psa3 G_018 | 2017 | PRJNA826129 | This study |
| Psa3 G_019 | 2017 | PRJNA826129 | This study |
| Psa3 G_020 | 2017 | PRJNA826129 | This study |
| Psa3 G_021 | 2017 | PRJNA826129 | This study |
| Psa3 G_036 | 2017 | PRJNA826129 | This study |
| Psa3 G_037 | 2017 | PRJNA826129 | This study |
| Psa3 G_039 | 2017 | PRJNA826129 | This study |
| Psa3 G_040 | 2017 | PRJNA826129 | This study |
| Psa3 G_041 | 2017 | PRJNA826129 | This study |
| Psa3 G_042 | 2017 | PRJNA826129 | This study |
| Psa3 G_043 | 2017 | PRJNA826129 | This study |
| Psa3 G_044 | 2017 | PRJNA826129 | This study |
| Psa3 G_045 | 2017 | PRJNA826129 | This study |
| Psa3 G_046 | 2017 | PRJNA826129 | This study |
| Psa3 G_047 | 2017 | PRJNA826129 | This study |
| Psa3 G_048 | 2017 | PRJNA826129 | This study |
| Psa3 G_050 | 2017 | PRJNA826129 | This study |
| Psa3 G_052 | 2017 | PRJNA826129 | This study |
| Psa3 G_058 | 2017 | PRJNA826129 | This study |
| Psa3 G_059 | 2017 | PRJNA826129 | This study |
| Psa3 G_060 | 2017 | PRJNA826129 | This study |
| Psa3 G_061 | 2017 | PRJNA826129 | This study |
| Psa3 G_062 | 2017 | PRJNA826129 | This study |
| Psa3 G_063 | 2017 | PRJNA826129 | This study |
| Psa3 G_064 | 2017 | PRJNA826129 | This study |
| Psa3 G_065 | 2017 | PRJNA826129 | This study |
| Psa3 G_066 | 2017 | PRJNA826129 | This study |
| Psa3 G_067 | 2017 | PRJNA826129 | This study |
| Psa3 G_068 | 2017 | PRJNA826129 | This study |
| Psa3 G_070 | 2017 | PRJNA826129 | This study |
| Psa3 G_071 | 2017 | PRJNA826129 | This study |
| Psa3 G_072 | 2017 | PRJNA826129 | This study |
| Psa3 G_073 | 2017 | PRJNA826129 | This study |
| Psa3 G_074 | 2017 | PRJNA826129 | This study |
| Psa3 G_075 | 2017 | PRJNA826129 | This study |
| Psa3 G_076 | 2017 | PRJNA826129 | This study |
| Psa3 G_077 | 2017 | PRJNA826129 | This study |
| Psa3 G_078 | 2017 | PRJNA826129 | This study |
| Psa3 G_079 | 2017 | PRJNA826129 | This study |
| Psa3 G_080 | 2017 | PRJNA826129 | This study |
| Psa3 G_081 | 2017 | PRJNA826129 | This study |
| Psa3 G_082 | 2017 | PRJNA826129 | This study |
| Psa3 G_083 | 2017 | PRJNA826129 | This study |
| Psa3 G_084 | 2017 | PRJNA826129 | This study |
| Psa3 G_085 | 2017 | PRJNA826129 | This study |
| Psa3 G_086 | 2017 | PRJNA826129 | This study |
| Psa3 G_087 | 2017 | PRJNA826129 | This study |
| Psa3 G_088 | 2017 | PRJNA826129 | This study |
| Psa3 H_010 | 2017 | PRJNA826129 | This study |
| Psa3 H_053 | 2017 | PRJNA826129 | This study |
| Psa3 H_054 | 2017 | PRJNA826129 | This study |
| Psa3 H_055 | 2017 | PRJNA826129 | This study |
| Psa3 H_056 | 2017 | PRJNA826129 | This study |
| Psa3 H_057 | 2017 | PRJNA826129 | This study |
| Psa3 H_089 | 2017 | PRJNA826129 | This study |
| Psa3 H_090 | 2017 | PRJNA826129 | This study |
| Psa3 H_091 | 2017 | PRJNA826129 | This study |
| Psa3 H_092 | 2017 | PRJNA826129 | This study |
| Psa3 H_093 | 2017 | PRJNA826129 | This study |
| Psa3 H_094 | 2017 | PRJNA826129 | This study |
| Psa3 H_095 | 2017 | PRJNA826129 | This study |
| Psa3 H_096 | 2017 | PRJNA826129 | This study |
| Psa3 HBL66A | 2017 | PRJNA527140 | University of Otago |
| Psa3 HL103 | 2017 | PRJNA527140 | University of Otago |
| Psa3 HL105 | 2017 | PRJNA527140 | University of Otago |
| Psa3 HL111 | 2017 | PRJNA527140 | University of Otago |
| Psa3 HL112 | 2017 | PRJNA527140 | University of Otago |
| Psa3 HL113 | 2017 | PRJNA527140 | University of Otago |
| Psa3 HL117 | 2017 | PRJNA527140 | University of Otago |
| Psa3 HL118 | 2017 | PRJNA527140 | University of Otago |
| Psa3 HL120 | 2017 | PRJNA527140 | University of Otago |
| Psa3 L121 | 2017 | PRJNA527140 | University of Otago |
| Psa3 L147 | 2017 | PRJNA527140 | University of Otago |
| Psa3 L206 | 2017 | PRJNA527140 | University of Otago |
| Psa3 L508 | 2017 | PRJNA527140 | University of Otago |
| Psa3 L60A | 2017 | PRJNA527140 | University of Otago |
| Psa3 L716B | 2017 | PRJNA527140 | University of Otago |
| Psa3 L72H | 2017 | PRJNA527140 | University of Otago |
| Psa3 n380 | 2017 | PRJNA527140 | University of Otago |
| Psa3 n386 | 2017 | PRJNA527140 | University of Otago |
| Psa3 n507 | 2017 | PRJNA527140 | University of Otago |
| Psa3 r929 | 2017 | PRJNA527140 | University of Otago |
| Psa3 r932 | 2017 | PRJNA527140 | University of Otago |
| Psa3 S12-1M | 2017 | PRJNA527140 | University of Otago |
| Psa3 S19 | 2017 | PRJNA527140 | University of Otago |
| Psa3 S7-1M | 2017 | PRJNA527140 | University of Otago |
| Psa3 SBC2 | 2017 | PRJNA527140 | University of Otago |
| Psa3 SBC7 | 2017 | PRJNA527140 | University of Otago |
| Psa3 SP1 | 2017 | PRJNA527140 | University of Otago |
| Psa3 SP11-7 | 2017 | PRJNA527140 | University of Otago |
| Psa3 SP13-1 | 2017 | PRJNA527140 | University of Otago |
| Psa3 SP14-1 | 2017 | PRJNA527140 | University of Otago |
| Psa3 SP16-1 | 2017 | PRJNA527140 | University of Otago |
| Psa3 SP18-1 | 2017 | PRJNA527140 | University of Otago |
| Psa3 SP20-1 | 2017 | PRJNA527140 | University of Otago |
| Psa3 SP21-1 | 2017 | PRJNA527140 | University of Otago |
| Psa3 SP22-1 | 2017 | PRJNA527140 | University of Otago |
| Psa3 SP23-2 | 2017 | PRJNA527140 | University of Otago |
| Psa3 SP24-2 | 2017 | PRJNA527140 | University of Otago |
| Psa3 SP26 | 2017 | PRJNA527140 | University of Otago |
| Psa3 SP27 | 2017 | PRJNA527140 | University of Otago |
| Psa3 SP29 | 2017 | PRJNA527140 | University of Otago |
| Psa3 SP37 | 2017 | PRJNA527140 | University of Otago |
| Psa3 SP38-1 | 2017 | PRJNA527140 | University of Otago |
| Psa3 SP43-1 | 2017 | PRJNA527140 | University of Otago |
| Psa3 SP44-2 | 2017 | PRJNA527140 | University of Otago |
| Psa3 G_097 | 2018 | PRJNA826132 | This study |
| Psa3 G_098 | 2018 | PRJNA826132 | This study |
| Psa3 G_099 | 2018 | PRJNA826132 | This study |
| Psa3 G_100 | 2018 | PRJNA826132 | This study |
| Psa3 G_101 | 2018 | PRJNA826132 | This study |
| Psa3 G_102 | 2018 | PRJNA826132 | This study |
| Psa3 G_103 | 2018 | PRJNA826132 | This study |
| Psa3 G_104 | 2018 | PRJNA826132 | This study |
| Psa3 G_105 | 2018 | PRJNA826132 | This study |
| Psa3 G_106 | 2018 | PRJNA826132 | This study |
| Psa3 G_107 | 2018 | PRJNA826132 | This study |
| Psa3 G_108 | 2018 | PRJNA826132 | This study |
| Psa3 G_109 | 2018 | PRJNA826132 | This study |
| Psa3 G_110 | 2018 | PRJNA826132 | This study |
| Psa3 G_111 | 2018 | PRJNA826132 | This study |
| Psa3 G_112 | 2018 | PRJNA826132 | This study |
| Psa3 G_118 | 2018 | PRJNA826132 | This study |
| Psa3 G_119 | 2018 | PRJNA826132 | This study |
| Psa3 G_120 | 2018 | PRJNA826132 | This study |
| Psa3 G_121 | 2018 | PRJNA826132 | This study |
| Psa3 G_123 | 2018 | PRJNA826132 | This study |
| Psa3 G_124 | 2018 | PRJNA826132 | This study |
| Psa3 G_125 | 2018 | PRJNA826132 | This study |
| Psa3 G_126 | 2018 | PRJNA826132 | This study |
| Psa3 G_127 | 2018 | PRJNA826132 | This study |
| Psa3 G_128 | 2018 | PRJNA826132 | This study |
| Psa3 G_129 | 2018 | PRJNA826132 | This study |
| Psa3 G_130 | 2018 | PRJNA826132 | This study |
| Psa3 G_131 | 2018 | PRJNA826132 | This study |
| Psa3 G_132 | 2018 | PRJNA826132 | This study |
| Psa3 G_133 | 2018 | PRJNA826132 | This study |
| Psa3 G_134 | 2018 | PRJNA826132 | This study |
| Psa3 G_135 | 2018 | PRJNA826132 | This study |
| Psa3 G_136 | 2018 | PRJNA826132 | This study |
| Psa3 G_137 | 2018 | PRJNA826132 | This study |
| Psa3 G_138 | 2018 | PRJNA826132 | This study |
| Psa3 G_139 | 2018 | PRJNA826132 | This study |
| Psa3 G_140 | 2018 | PRJNA826132 | This study |
| Psa3 G_141 | 2018 | PRJNA826132 | This study |
| Psa3 G_179 | 2018 | PRJNA826132 | This study |
| Psa3 G_180 | 2018 | PRJNA826132 | This study |
| Psa3 G_181 | 2018 | PRJNA826132 | This study |
| Psa3 G_182 | 2018 | PRJNA826132 | This study |
| Psa3 G_183 | 2018 | PRJNA826132 | This study |
| Psa3 G_184 | 2018 | PRJNA826132 | This study |
| Psa3 G_185 | 2018 | PRJNA826132 | This study |
| Psa3 G_186 | 2018 | PRJNA826132 | This study |
| Psa3 G_187 | 2018 | PRJNA826132 | This study |
| Psa3 G_188 | 2018 | PRJNA826132 | This study |
| Psa3 G_189 | 2018 | PRJNA826132 | This study |
| Psa3 G_190 | 2018 | PRJNA826132 | This study |
| Psa3 G_191 | 2018 | PRJNA826132 | This study |
| Psa3 G_192 | 2018 | PRJNA826132 | This study |
| Psa3 H_113 | 2018 | PRJNA826132 | This study |
| Psa3 H_114 | 2018 | PRJNA826132 | This study |
| Psa3 H_115 | 2018 | PRJNA826132 | This study |
| Psa3 H_116 | 2018 | PRJNA826132 | This study |
| Psa3 H_142 | 2018 | PRJNA826132 | This study |
| Psa3 H_143 | 2018 | PRJNA826132 | This study |
| Psa3 H_144 | 2018 | PRJNA826132 | This study |
| Psa3 H_145 | 2018 | PRJNA826132 | This study |
| Psa3 H_146 | 2018 | PRJNA826132 | This study |
| Psa3 H_147 | 2018 | PRJNA826132 | This study |
| Psa3 H_193 | 2018 | PRJNA826132 | This study |
| Psa3 H_194 | 2018 | PRJNA826132 | This study |
| Psa3 H_195 | 2018 | PRJNA826132 | This study |
| Psa3 H_196 | 2018 | PRJNA826132 | This study |
| Psa3 H_197 | 2018 | PRJNA826132 | This study |
| Psa3 B_287 | 2020 | PRJNA826143 | This study |
| Psa3 B_288 | 2020 | PRJNA826143 | This study |
| Psa3 B_289 | 2020 | PRJNA826143 | This study |
| Psa3 B_290 | 2020 | PRJNA826143 | This study |
| Psa3 B_299 | 2020 | PRJNA826143 | This study |
| Psa3 B_300 | 2020 | PRJNA826143 | This study |
| Psa3 B_301 | 2020 | PRJNA826143 | This study |
| Psa3 B_439 | 2020 | PRJNA826143 | This study |
| Psa3 B_440 | 2020 | PRJNA826143 | This study |
| Psa3 G_291 | 2020 | PRJNA826143 | This study |
| Psa3 G_292 | 2020 | PRJNA826143 | This study |
| Psa3 G_293 | 2020 | PRJNA826143 | This study |
| Psa3 G_302 | 2020 | PRJNA826143 | This study |
| Psa3 G_303 | 2020 | PRJNA826143 | This study |
| Psa3 G_304 | 2020 | PRJNA826143 | This study |
| Psa3 G_305 | 2020 | PRJNA826143 | This study |
| Psa3 G_306 | 2020 | PRJNA826143 | This study |
| Psa3 G_307 | 2020 | PRJNA826143 | This study |
| Psa3 G_308 | 2020 | PRJNA826143 | This study |
| Psa3 G_309 | 2020 | PRJNA826143 | This study |
| Psa3 G_310 | 2020 | PRJNA826143 | This study |
| Psa3 G_311 | 2020 | PRJNA826143 | This study |
| Psa3 G_312 | 2020 | PRJNA826143 | This study |
| Psa3 G_313 | 2020 | PRJNA826143 | This study |
| Psa3 G_314 | 2020 | PRJNA826143 | This study |
| Psa3 G_315 | 2020 | PRJNA826143 | This study |
| Psa3 G_316 | 2020 | PRJNA826143 | This study |
| Psa3 G_317 | 2020 | PRJNA826143 | This study |
| Psa3 G_318 | 2020 | PRJNA826143 | This study |
| Psa3 G_319 | 2020 | PRJNA826143 | This study |
| Psa3 G_320 | 2020 | PRJNA826143 | This study |
| Psa3 G_321 | 2020 | PRJNA826143 | This study |
| Psa3 G_322 | 2020 | PRJNA826143 | This study |
| Psa3 G_356 | 2020 | PRJNA826143 | This study |
| Psa3 G_357 | 2020 | PRJNA826143 | This study |
| Psa3 G_358 | 2020 | PRJNA826143 | This study |
| Psa3 G_359 | 2020 | PRJNA826143 | This study |
| Psa3 G_360 | 2020 | PRJNA826143 | This study |
| Psa3 G_361 | 2020 | PRJNA826143 | This study |
| Psa3 G_409 | 2020 | PRJNA826143 | This study |
| Psa3 G_410 | 2020 | PRJNA826143 | This study |
| Psa3 G_411 | 2020 | PRJNA826143 | This study |
| Psa3 G_412 | 2020 | PRJNA826143 | This study |
| Psa3 G_413 | 2020 | PRJNA826143 | This study |
| Psa3 G_414 | 2020 | PRJNA826143 | This study |
| Psa3 G_415 | 2020 | PRJNA826143 | This study |
| Psa3 G_416 | 2020 | PRJNA826143 | This study |
| Psa3 G_417 | 2020 | PRJNA826143 | This study |
| Psa3 G_418 | 2020 | PRJNA826143 | This study |
| Psa3 G_419 | 2020 | PRJNA826143 | This study |
| Psa3 G_420 | 2020 | PRJNA826143 | This study |
| Psa3 G_421 | 2020 | PRJNA826143 | This study |
| Psa3 G_422 | 2020 | PRJNA826143 | This study |
| Psa3 G_423 | 2020 | PRJNA826143 | This study |
| Psa3 G_424 | 2020 | PRJNA826143 | This study |
| Psa3 G_434 | 2020 | PRJNA826143 | This study |
| Psa3 G_435 | 2020 | PRJNA826143 | This study |
| Psa3 G_441 | 2020 | PRJNA826143 | This study |
| Psa3 G_442 | 2020 | PRJNA826143 | This study |
| Psa3 G_443 | 2020 | PRJNA826143 | This study |
| Psa3 G_444 | 2020 | PRJNA826143 | This study |
| Psa3 G_445 | 2020 | PRJNA826143 | This study |
| Psa3 G_446 | 2020 | PRJNA826143 | This study |
| Psa3 G_447 | 2020 | PRJNA826143 | This study |
| Psa3 G_448 | 2020 | PRJNA826143 | This study |
| Psa3 G_449 | 2020 | PRJNA826143 | This study |
| Psa3 G_450 | 2020 | PRJNA826143 | This study |
| Psa3 G_451 | 2020 | PRJNA826143 | This study |
| Psa3 H_294 | 2020 | PRJNA826143 | This study |
| Psa3 H_295 | 2020 | PRJNA826143 | This study |
| Psa3 H_324 | 2020 | PRJNA826143 | This study |
| Psa3 H_325 | 2020 | PRJNA826143 | This study |
| Psa3 H_326 | 2020 | PRJNA826143 | This study |
| Psa3 H_327 | 2020 | PRJNA826143 | This study |
| Psa3 H_328 | 2020 | PRJNA826143 | This study |
| Psa3 H_329 | 2020 | PRJNA826143 | This study |
| Psa3 H_330 | 2020 | PRJNA826143 | This study |
| Psa3 H_344 | 2020 | PRJNA826143 | This study |
| Psa3 H_345 | 2020 | PRJNA826143 | This study |
| Psa3 H_362 | 2020 | PRJNA826143 | This study |
| Psa3 H_363 | 2020 | PRJNA826143 | This study |
| Psa3 H_364 | 2020 | PRJNA826143 | This study |
| Psa3 H_365 | 2020 | PRJNA826143 | This study |
| Psa3 H_366 | 2020 | PRJNA826143 | This study |
| Psa3 H_367 | 2020 | PRJNA826143 | This study |
| Psa3 H_368 | 2020 | PRJNA826143 | This study |
| Psa3 H_369 | 2020 | PRJNA826143 | This study |
| Psa3 H_370 | 2020 | PRJNA826143 | This study |
| Psa3 H_371 | 2020 | PRJNA826143 | This study |
| Psa3 H_372 | 2020 | PRJNA826143 | This study |
| Psa3 H_373 | 2020 | PRJNA826143 | This study |
| Psa3 H_375 | 2020 | PRJNA826143 | This study |
| Psa3 H_376 | 2020 | PRJNA826143 | This study |
| Psa3 H_377 | 2020 | PRJNA826143 | This study |
| Psa3 H_378 | 2020 | PRJNA826143 | This study |
| Psa3 H_379 | 2020 | PRJNA826143 | This study |
| Psa3 H_380 | 2020 | PRJNA826143 | This study |
| Psa3 H_381 | 2020 | PRJNA826143 | This study |
| Psa3 H_382 | 2020 | PRJNA826143 | This study |
| Psa3 H_383 | 2020 | PRJNA826143 | This study |
| Psa3 H_384 | 2020 | PRJNA826143 | This study |
| Psa3 H_385 | 2020 | PRJNA826143 | This study |
| Psa3 H_386 | 2020 | PRJNA826143 | This study |
| Psa3 H_387 | 2020 | PRJNA826143 | This study |
| Psa3 H_388 | 2020 | PRJNA826143 | This study |
| Psa3 H_389 | 2020 | PRJNA826143 | This study |
| Psa3 H_390 | 2020 | PRJNA826143 | This study |
| Psa3 H_391 | 2020 | PRJNA826143 | This study |
| Psa3 H_392 | 2020 | PRJNA826143 | This study |
| Psa3 H_393 | 2020 | PRJNA826143 | This study |
| Psa3 H_394 | 2020 | PRJNA826143 | This study |
| Psa3 H_395 | 2020 | PRJNA826143 | This study |
| Psa3 H_396 | 2020 | PRJNA826143 | This study |
| Psa3 H_397 | 2020 | PRJNA826143 | This study |
| Psa3 H_398 | 2020 | PRJNA826143 | This study |
| Psa3 H_399 | 2020 | PRJNA826143 | This study |
| Psa3 H_400 | 2020 | PRJNA826143 | This study |
| Psa3 H_401 | 2020 | PRJNA826143 | This study |
| Psa3 H_402 | 2020 | PRJNA826143 | This study |
| Psa3 H_403 | 2020 | PRJNA826143 | This study |
| Psa3 H_404 | 2020 | PRJNA826143 | This study |
| Psa3 H_405 | 2020 | PRJNA826143 | This study |
| Psa3 H_406 | 2020 | PRJNA826143 | This study |
| Psa3 H_407 | 2020 | PRJNA826143 | This study |
| Psa3 H_408 | 2020 | PRJNA826143 | This study |
| Psa3 H_425 | 2020 | PRJNA826143 | This study |
| Psa3 H_426 | 2020 | PRJNA826143 | This study |
| Psa3 H_427 | 2020 | PRJNA826143 | This study |
| Psa3 H_428 | 2020 | PRJNA826143 | This study |
| Psa3 H_429 | 2020 | PRJNA826143 | This study |
| Psa3 H_430 | 2020 | PRJNA826143 | This study |
| Psa3 H_431 | 2020 | PRJNA826143 | This study |
| Psa3 H_432 | 2020 | PRJNA826143 | This study |
| Psa3 H_433 | 2020 | PRJNA826143 | This study |
| Psa3 H_436 | 2020 | PRJNA826143 | This study |
| Psa3 H_437 | 2020 | PRJNA826143 | This study |
| Psa3 H_438 | 2020 | PRJNA826143 | This study |
| Psa3 H_452 | 2020 | PRJNA826143 | This study |
| Psa3 H_453 | 2020 | PRJNA826143 | This study |
| Psa3 H_454 | 2020 | PRJNA826143 | This study |
| Psa3 H_455 | 2020 | PRJNA826143 | This study |
| Psa3 H_456 | 2020 | PRJNA826143 | This study |
| Psa3 H_457 | 2020 | PRJNA826143 | This study |
| Psa3 H_458 | 2020 | PRJNA826143 | This study |
| Psa3 H_459 | 2020 | PRJNA826143 | This study |
| Psa3 H_460 | 2020 | PRJNA826143 | This study |
| Psa3 N_296 | 2020 | PRJNA826143 | This study |
| Psa3 N_297 | 2020 | PRJNA826143 | This study |
| Psa3 N_298 | 2020 | PRJNA826143 | This study |
| Psa3 G_486 | 2022 | PRJNA1165295 | This study |
| Psa3 G_487 | 2022 | PRJNA1165295 | This study |
| Psa3 G_488 | 2022 | PRJNA1165295 | This study |
| Psa3 G_490 | 2022 | PRJNA1165295 | This study |
| Psa3 G_495 | 2022 | PRJNA1165295 | This study |
| Psa3 G_499 | 2022 | PRJNA1165295 | This study |
| Psa3 G_500 | 2022 | PRJNA1165295 | This study |
| Psa3 G_503 | 2022 | PRJNA1165295 | This study |
| Psa3 G_508 | 2022 | PRJNA1165295 | This study |
| Psa3 G_509 | 2022 | PRJNA1165295 | This study |
| Psa3 G_511 | 2022 | PRJNA1165295 | This study |
| Psa3 G_514 | 2022 | PRJNA1165295 | This study |
| Psa3 G_515 | 2022 | PRJNA1165295 | This study |
| Psa3 G_517 | 2022 | PRJNA1165295 | This study |
| Psa3 G_518 | 2022 | PRJNA1165295 | This study |
| Psa3 G_520 | 2022 | PRJNA1165295 | This study |
| Psa3 G_523 | 2022 | PRJNA1165295 | This study |
| Psa3 G_526 | 2022 | PRJNA1165295 | This study |
| Psa3 G_527 | 2022 | PRJNA1165295 | This study |
| Psa3 G_528 | 2022 | PRJNA1165295 | This study |
| Psa3 G_529 | 2022 | PRJNA1165295 | This study |
| Psa3 G_531 | 2022 | PRJNA1165295 | This study |
| Psa3 G_533 | 2022 | PRJNA1165295 | This study |
| Psa3 H_489 | 2022 | PRJNA1165295 | This study |
| Psa3 H_491 | 2022 | PRJNA1165295 | This study |
| Psa3 H_492 | 2022 | PRJNA1165295 | This study |
| Psa3 H_493 | 2022 | PRJNA1165295 | This study |
| Psa3 H_494 | 2022 | PRJNA1165295 | This study |
| Psa3 H_496 | 2022 | PRJNA1165295 | This study |
| Psa3 H_497 | 2022 | PRJNA1165295 | This study |
| Psa3 H_498 | 2022 | PRJNA1165295 | This study |
| Psa3 H_501 | 2022 | PRJNA1165295 | This study |
| Psa3 H_502 | 2022 | PRJNA1165295 | This study |
| Psa3 H_504 | 2022 | PRJNA1165295 | This study |
| Psa3 H_505 | 2022 | PRJNA1165295 | This study |
| Psa3 H_506 | 2022 | PRJNA1165295 | This study |
| Psa3 H_507 | 2022 | PRJNA1165295 | This study |
| Psa3 H_510 | 2022 | PRJNA1165295 | This study |
| Psa3 H_512 | 2022 | PRJNA1165295 | This study |
| Psa3 H_513 | 2022 | PRJNA1165295 | This study |
| Psa3 H_516 | 2022 | PRJNA1165295 | This study |
| Psa3 H_519 | 2022 | PRJNA1165295 | This study |
| Psa3 H_521 | 2022 | PRJNA1165295 | This study |
| Psa3 H_522 | 2022 | PRJNA1165295 | This study |
| Psa3 H_524 | 2022 | PRJNA1165295 | This study |
| Psa3 H_525 | 2022 | PRJNA1165295 | This study |
| Psa3 H_530 | 2022 | PRJNA1165295 | This study |
